# Novel Fusarium Wilt Resistance Genes Uncovered in the Wild Progenitors and Heirloom Cultivars of Strawberry

**DOI:** 10.1101/2021.12.07.471687

**Authors:** Dominique D. A. Pincot, Mitchell J. Feldmann, Michael A. Hardigan, Mishi V. Vachev, Peter M. Henry, Thomas R. Gordon, Alan Rodriguez, Nicolas Cobo, Glenn S. Cole, Gitta L. Coaker, Steven J. Knapp

## Abstract

Fusarium wilt, a soilborne disease caused by *Fusarium oxysporum* f. sp. *fragariae*, poses a significant threat to strawberry (*Fragaria* × *ananassa*) production in many parts of the world. This pathogen causes wilting, collapse, and death in susceptible genotypes. We previously identified a dominant gene (*FW1*) on chromosome 2B that confers resistance to race 1 of the pathogen and hypothesized that gene-for-gene resistance to Fusarium wilt was widespread in strawberry. To explore this, a genetically diverse collection of heirloom and modern cultivars and wild octoploid ecotypes were screened for resistance to Fusarium wilt races 1 and 2. Here we show that resistance to both races is widespread and that resistance to race 1 is mediated by dominant genes (*FW1, FW2, FW3, FW4*, and *FW5*) on three non-homoeologous chromosomes (1A, 2B, and 6B). The resistance proteins encoded by these genes are not yet known; however, plausible candidates were identified that encode pattern recognition receptor or other proteins known to mediate gene-for-gene resistance in plants.

High-throughput genotyping assays for SNPs in linkage disequilibrium with *FW1*-*FW5* were developed to facilitate marker-assisted selection and accelerate the development of race 1 resistant cultivars. This study laid the foundation for identifying the genes encoded by *FW1-FW5*, in addition to exploring the genetics of resistance to race 2 and other races of the pathogen, as a precaution to averting a Fusarium wilt pandemic.

**Key Message:** **Several race-specific resistance genes were identified and rapidly deployed via marker-assisted selection to develop strawberry cultivars resistant to Fusarium wilt, a devastating soil-borne disease**.

## Introduction

*Fusarium oxysporum*, a widespread soil-borne pathogen, causes vascular wilt disease in several economically important plants (Michielse and Rep, 2009; Dean et al., 2012), in addition to the broad spectrum human disease known as ‘fusariosis’ (Dignani and Anaissie, 2004; Nucci and Anaissie, 2007; Batista et al., 2020). *F. oxysporum* is one of the most destructive plant-pathogenic fungi worldwide, with a long and storied history of outbreaks and epidemics that have caused significant production losses and disrupted food and fiber production (Dean et al., 2012). One of the earliest reports of the disease arose from outbreaks on banana (*Musa acuminata* Colla) in the late 1800s that progressively annihilated the widely grown susceptible cultivar ‘Gros Michel’, forced the abandonment of export plantations, and caused a gradual, albeit inexorable shift in production from susceptible ‘Gros Michel’ to resistant ‘Cavendish’ cultivars (Ploetz, 2015; Dale et al., 2017; Pegg et al., 2019). Similar production shifts have unfolded over the last century in tomato (*Solanum lycopersicum* L.), cotton (*Gossypium hirsutum* L.), and other economically important plants (Michielse and Rep, 2009), and more recently strawberry (*Fragaria* × *ananassa* Duchesne ex Rozier) (Pincot et al., 2018; Henry et al., 2021). The discovery of sources of resistance and development and deployment of resistant cultivars has been critical for limiting disease losses and sustaining agricultural production in strawberry and other host plants affected by the pathogen (Dean et al., 2012; Gordon, 2017; Pincot et al., 2018; Henry et al., 2019).

Fusarium wilt of strawberry is caused by *F. oxysporum* f. sp. *fragariae* (Fof), one of more than 100 documented host-specific pathogens (formae speciales), many of which have been widely disseminated (Gordon, 2017). Although the strawberry-specific Fof has been reported in many countries, the disease has received the most attention in Japan, South Korea, Australia, and California, between which virulent strains have been disseminated (Gordon, 2017; Henry et al., 2017, 2021). Fusarium wilt was first reported on strawberry in Australia in the 1960s (Winks and Williams, 1965), and was not reported on strawberry in California until the mid-2000s (Koike et al., 2009; Koike and Gordon, 2015). The disease has been aggressively spreading and poses a serious threat to production in California (Koike and Gordon, 2015; Henry et al., 2017, 2019).

Fusarium wilt has not yet become a serious threat to production everywhere strawberries are grown; however, there is a significant risk of virulent strains being disseminated through global trade, and the ever present danger of the evolution and emergence of virulent races of the pathogen that defeat known resistance (*R*) genes (Henry et al., 2021). One of the motivations for the present study was to prepare for that inevitability by delving more deeply into the genetics of resistance and developing the resources and knowledge needed to accelerate the development of Fusarium wilt resistant cultivars through marker-assisted selection (MAS). To that end, we initiated studies in 2015 to identify sources of resistance to California isolates of the pathogen and shed light on the genetics of resistance to Fusarium wilt in strawberry (Pincot et al., 2018). The prevalence, diversity, strength of resistance, and genetic mechanisms underlying resistance to Fusarium wilt were unknown when those studies were initiated (Mori et al., 2005; Paynter et al., 2014; Pincot et al., 2018). Significant insights into the *Fragaria-Fusarium* pathosystem have since emerged.

Pincot et al. (2018) identified multiple sources of resistance to Fusarium wilt in a closed breeding population developed at the University of California, Davis (hereafter designated as the ‘California’ population). The isolate they used (AMP132) was subsequently classified as Fof race 1 (Henry et al., 2021). From the resistance phenotypes of plants artificially inoculated with AMP132, they observed a nearly bimodal distribution of resistant and susceptible individuals in a genome-wide association study (GWAS) of the California population, observed near-Mendelian distributions for resistance phenotypes in segregating populations, and showed that resistance to AMP132 was conferred by a single dominant gene (*FW1*) in the California population. The resistant allele (*FW1*) had a low frequency (0.16) and was only homozygous in 3% of the resistant individuals in the California population (Pincot et al., 2018). From analyses of pedigree records and haplotypes of SNP markers in linkage disequilibrium with the *FW1* locus, Pincot et al. (2018) predicted that 99% of the resistant individuals in the California population carried *FW1*. They concluded that the resistant allele (*FW1*) had fortuitously survived early breeding bottlenecks and originated in the earliest known ancestors of the California population (Pincot et al., 2018; Hardigan et al., 2021b; Pincot et al., 2021).

Pincot et al. (2018) screened two non-California cultivars (Guardian and Earliglow), both of which were shown to be resistant to race 1 and had SNP marker haplotypes different from the *FW1* SNP marker haplotype. The only AMP132-resistant cultivar in the California population without the *FW1* SNP marker haplotype was the heirloom cultivar ‘Wiltguard’. We speculated that Earliglow, Guardian, and Wiltguard might carry novel *R*-genes, a hypothesis tested in the present study. To build on earlier findings in the California population and develop a deeper understanding of the genetics of resistance, we screened a diverse collection of elite and exotic germplasm accessions (clonally preserved individuals) for resistance to race 1 and selected several additional race 1 resistant donors for further study. Here we show that resistance to race 1 is widespread in elite and exotic germplasm, including geographically diverse ecotypes of the wild octoploid progenitors of strawberry (*F. chiloensis* and *F. virginiana*).

Plant genes that confer strong race-specific resistance frequently encode proteins with nucleotide-binding leucinerich repeat domains (NLRs) or surface localized pattern recognition receptors (PRRs) (Lolle et al., 2020). Several of the previously described Fusarium wilt *R*-genes encode proteins with NLR and PRR architecture (Ori et al., 1997; Joobeur et al., 2004; Diener and Ausubel, 2005; Michielse and Rep, 2009; Lv et al., 2014; Catanzariti et al., 2015; Gonzalez-Cendales et al., 2016; Catanzariti et al., 2017). *R*-genes that confer resistance to *F. oxysporum* f. sp. *lycopersici* in tomato (*I, I-2, I-3, I-4*, and *I-7*) are among the most well studied examples (Sela-Buurlage et al., 2001; Hemming et al., 2004; Houterman et al., 2009; Michielse and Rep, 2009). PRRs are capable of recognizing conserved pathogen features and extracellular effectors, while NLR receptors recognize secreted pathogen effectors inside plant cells, resulting in disease resistance (Jones et al., 2016; Albert et al., 2020; Lolle et al., 2020). Although the gene encoded by *FW1* has not yet been identified, we posited that *FW1* might encode an NLR or PRR immune receptor protein that recognizes an effector protein encoded by Fof race 1 isolates (called AvrFW1).

Because *R*-genes often have short-lived utility (Mundt, 2014), the continual discovery and deployment of novel *R*-genes has been critical for keeping pace with the evolution of pathogen races in the gene-for-gene ‘arms race’ (Hammond-Kosack and Jones, 1996, 1997; Boller and He, 2009; Dangl et al., 2013; Chiang and Coaker, 2015). The durability of *FW1* and other race-specific *R*-genes is uncertain (Mundt, 2014, 2018), and depends on the speed of emergence of novel Fof races through pathogen mutation (White et al., 2000; Rouxel and Balesdent, 2010; Henry et al., 2021). If *FW1* encodes an NLR or PRR, a mutation of *AvrFW1* could lead to an evasion of host immune perception and regained pathogenicity (Zhang and Coaker, 2017). Currently, only race 1 isolates of Fof have been found in California, and none cause disease in cultivars carrying the dominant *FW1* allele (Henry et al., 2017; Pincot et al., 2018; Henry et al., 2021). However, race 2 isolates that cause disease on *FW1*-carrying cultivars have been observed (Henry et al., 2021). The identification of race 2 reinforces the expectation that novel strains of the pathogen could eventually evolve and defeat defeat race 1 *R*-genes through mutation, loss, or expression polymorphism in *AvrFW1*.

The identification of *FW1* and *AvrFW1* and advances in the development of genomic resources for *Fragaria* and *Fusarium* laid the foundation for the present study. *FW1* was originally discovered by GWAS using a diploid reference genome (Pincot et al., 2018). The approximate location of *FW1* in the octoploid genome was subsequently ascertained by genetic mapping in octoploid segregating populations genotyped with a single nucleotide polymorphsim (SNP) array designed with probe DNA sequences anchored to a diploid reference genome (Bassil et al., 2015; Verma et al., 2017; Pincot et al., 2018). The octoploid genome has since been sequenced (Edger et al., 2019; Hardigan et al., 2021a), thereby opening the way for octoploid genome-informed breeding and genetic studies in strawberry. Those genome assemblies supplied the foundation for several additional technical advances, the most important of which were the genome-wide discovery and physical and genetic mapping of millions of DNA variants in the octoploid genome, the development of 50K and 850K SNP genotyping arrays with probe DNA sequences uniformly distributed and anchored to physical positions throughout the octoploid genome, and telomere-to-telomere resolution of the A, B, C, and D subgenomes of octoploid strawberry (Hardigan et al., 2020, 2021b). These breakthroughs and resources were critical for the present study, which included: (a) pinpointing the genomic location of the *FW1* locus and four newly discovered Fusarium wilt resistance loci (*FW2, FW3, FW4*, and *FW5*); (b) expanding the database of octoploid germplasm accessions screened for resistance to Fusarium wilt races 1 and 2; (c) identifying SNPs and other DNA variants in linkage disequilibrium with *FW1*-*FW5*; and (d) identifying plausible candidate genes for *FW1*-*FW5* through genotype-to-phenotype associations. Finally, we describe high-throughput genotyping assays for SNPs in strong LD with *FW1*-*FW5* to facilitate the development of Fusarium wilt resistant cultivars through MAS.

## Materials and Methods

### Plant Material

The plant materials for our studies included 309 *F*. × *ananassa*, 62 *F. chiloensis*, and 40 *F. virginiana* germplasm accessions (individuals) preserved in the University of California, Davis (UC Davis) Strawberry Germplasm Collection or the United States Department of Agriculture, Agricultural Research Service, National Plant Germplasm System (USDA-ARS NPGS), National Clonal Germplasm Repository, Corvallis, Oregon (https://www.ars-grin.gov/). The original ‘mother’ plants of individuals acquired from the USDA were asexually multiplied in a Winters, CA field nursery and preserved in the UC Davis Strawberry Germplasm Collection throughout the course of our studies (Online Resource 1). Bare-root plants (clones) of every individual were produced by asexual multiplication in high-elevation (1,294 m) field nurseries in Dorris, CA from mother plants propagated in low-elevation (41 m) field nurseries in Winters, CA. The mother plants were planted mid-April and daughter plants were harvested and trimmed in mid-October and stored in plastic bags at 3.5°C for two to three weeks before pathogen inoculation and planting. The daughter plants for growth chamber and greenhouse experiments were stored at -2.2°C for 5 to 27 weeks and ultimately thawed and stored at 3.5°C for one to three days prior to pathogen inoculation and planting.

S_1_ families were developed by self-pollinating three Fusarium wilt race 1 resistant *F*. × *ananassa* cultivars identified by Pincot et al. (2018): Guardian (PI551407), Wiltguard (PI551669; 52C016P007), and Earliglow (PI551394). An S_1_ family was developed by self-pollinating a resistant individual (17C327P010) we identified in a population developed by crossing the susceptible cultivar Cabrillo with the resistant *F*. *virginiana* subsp. *glauca* ecotype PI612500. An S_2_ family was developed by self-pollinating 61S016P006, a highly resistant S_1_ individual identified in our resistance screening study. These individuals were known from genome-wide DNA profiling to be highly heterozygous and predicted *a priori* to either be heterozygous or homozygous for alleles affecting resistance. We developed interspecific full-sib families by crossing a susceptible *F. × ananassa* parent (12C089P002) with race 1 resistant ecotypes of *Fragaria virginiana* subsp. *virginiana* (PI552277), *Fragaria chiloensis* subsp. *patagonica* (PI602575), and *Fragaria virginiana* subsp. *grayana* (PI612569). These ecotypes were identified in the present study, known to be highly heterozygous from genome-wide DNA profiling, and, as before, predicted *a priori* to either be heterozygous or homozygous for alleles affecting resistance. The parents of these populations were grown in greenhouses at the UC Davis. S_1_ and S_2_ family seeds were produced by hand pollinating unemasculated flowers of Guardian, Wiltguard, Earliglow, 17C327P010, and 61S016P006. The PI612569 × 12C089P002, 12C089P002 × PI602575, and PI552277 × 12C089P002 full-sib families were produced by emasculating flowers on greenhouse grown plants of the female parent and hand pollinating the emasculated flowers with pollen from male parents. Ripe fruit were harvested and macerated in a pectinase solution (0.6 g/L) to separate achenes (seeds) from receptacles. Seeds were scarified by soaking in a 36 normal sulfuric acid solution for 16 min. Scarified seeds were germinated on moistened blotter paper at room temperature (approximately 22-24°C). Seedlings were transplanted to sterilized soil and were greenhouse grown for 9 months in Winters, CA before transplanting to the field, or were grown in a growth chamber for two to four months in Davis, CA before transplanting to the greenhouse.

### Artificial Inoculation Protocols and Disease Resistance Phenotyping

The plants for our experiments were artificially inoculated with race 1 (AMP132) and 2 (MAFF727510) isolates of *F. oxysporum* f. sp. *fragariae* using previously described protocols (Pincot et al., 2018; Henry et al., 2021). The AMP132 isolate originated in California, whereas the MAFF727510 isolate originated in Japan (Gordon et al., 2016; Henry et al., 2017; Pincot et al., 2018; Henry et al., 2021). To produce spores, the pathogen was grown on potato dextrose agar or Kerr’s broth under continuous fluorescent lighting at room temperature, as previously described (Pincot et al., 2018; Henry et al., 2021). Crude suspensions were passed through two layers of sterilized cheesecloth to remove hyphae. Spore densities were estimated using a haemocytometer and diluted with either sterile DI water (AMP132) or 0.1% water agar (MAFF727510) to a final density of 5 × 10^6^ spores/ml. Seedling and bare-root plants were inoculated by submerging their root systems up to the crown in the spore suspension for 7-8 min. prior to planting.

The individuals in these studies were visually phenotyped for resistance to Fusarium wilt over multiple post-inoculation time points using an ordinal disease rating scale from 1 (highly resistant) to 5 (highly susceptible) (Gordon et al., 2016; Henry et al., 2017; Pincot et al., 2018; Henry et al., 2021). For our field studies, individual plants were phenotyped once per week for four to eight consecutive weeks beginning in early June. Symptoms were observed on plants 26- to 36-weeks post-inoculation. For greenhouse and growth chamber studies, entries were phenotyped weekly for six to 12 weeks post-inoculation. For field, greenhouse, and growth chamber experiments, the onset and progression of disease symptoms among resistant and susceptible checks were used as guides for initiating and terminating phenotyping.

### Race 1 Resistance Screening Experiments

Our race 1 resistance screening experiments were conducted over a three year period (2016-17 to 2018-19) at the UC Davis Plant Pathology Farm. The plants for these experiments were artificially inoculated with the AMP132 isolate of the pathogen. Strawberries had not been previously grown in the fields selected for our studies. The fields were tilled and disked prior to fumigation and were broadcast-fumigated in October of each year with a 60:40 mixture of chloropicrin:1,3-dichloropropene (Pic-Clor 60, Cardinal Professional Products, Woodland, CA) at 560.4 kg/ha. The entire field was sealed with an impermeable plastic film for one-week post-fumigation before shaping 15.3 cm tall × 76.2 cm center-to-center raised beds. Sub-surface irrigation drip tape was installed longitudinally along the beds followed by black plastic mulch with a single row of planting holes spaced 30.5 cm apart. Artificially inoculated plants were transplanted in mid-November both years. The fields were fertilized with approximately 198 kg/ha of nitrogen over the growing season and irrigated as needed to prevent water stress.

For the 2016-17 field experiment, 344 germplasm accessions (identified in Online Resource 1) were screened for resistance to AMP132 and were part of a study that included 565 germplasm accessions developed at UC Davis, which is hereafter identified as the ‘California’ population. The resistance phenotypes for the latter were previously reported by Pincot et al. (2018). Collectively, 981 germplasm accessions were screened in the 2016-17 field study. These were arranged in a square lattice experiment design with four single-plant replicates per entry (Hinkelmann and Kempthorne, 1994). The experiment design and randomizations of entries within incomplete blocks were generated with the R package *agricolae* (De Mendiburu, 2015). For the 2017-18 and 2018-19 field experiments, 144 ‘host differential panel’ individuals were screened for resistance to AMP132 (identified in Online Resource 1). These individuals were arranged in a 12 *times* 12 square lattice experiment design with four single-plant replicates per entry as described above.

Guardian, Wiltguard, and Earliglow S_1_ and 61S016P006 S_2_ populations were screened for resistance to AMP132 in the 2016-17 field study. Ninety-nine Guardian S_1_ and 98 Wiltguard S_1_ individuals were phenotyped and genotyped and 85 Earliglow S_1_ and 77 61S016P006 *S*_2_ individuals were phenotyped. Nine-month-old S_1_ or S_2_ plants started as seedlings and asexually multiplied bare-root plants of the parents were artificially inoculated with AMP132, transplanted to the field in March 2018, and visually phenotyped weekly for six to 11 weeks post-inoculation.

The 12C089P002 × PI602575 (*n* = 76), PI552277 × 12C089P002 (*n* = 111), and PI612569 × 12C089P002 (*n* = 83) full-sib families, 17C327P010 S_1_ (*n* = 126) family, and parents of these families were screened for resistance to AMP132 in greenhouse experiments at UC Davis. Two to four-month-old seedlings of the progeny and bare-root plants of the parents were artificially inoculated with AMP132 and planted in February 2019 (17C327P010 S_1_), June 2019 (PI552277 × 12C089P002, PI612569 × 12C089P002), or November 2019 (12C089P002 × PI602575) into 10.2 ×10.2 × 15.2 cm plastic pots filled with 3 parts coir : 1 part perlite and phenotyped weekly for six to 12 weeks post-inoculation. Four uninoculated and four inoculated single-plant replicates of the parents were arranged in completely randomized experiment designs. The plants were irrigated with a dilute nutrient solution as needed to maintain adequate soil moisture. The 12C089P002 × PI602575 and PI552277 × 12C089P002 populations were genotyped with a 50K Axiom SNP array (Hardigan et al., 2020).

### Race 2 Resistance Screening Experiments

We screened the host differential panel (*n* = 144 individuals) for resistance to the MAFF727510 isolate of Fof race 2 in a growth chamber at the UC Davis Controlled Environment Facility in 2018-19. Two single-plant replicates/individual were arranged in a randomized complete block experiment design. The entire experiment was repeated twice, resulting in four clonal replications/individual. The bare-root plants for these experiments were produced in high-elevation nurseries, preserved in cold storage, artificially inoculated with the MAFF727510 isolate, transplanted into 10.2 ×10.2 × 15.2 cm plastic pots filled with a 4 parts sphagnum peat moss : 1 part perlite (Sunshine Mix #1; Sun Gro Horticulture, Agawam, MA), and phenotyped weekly for six to 12 weeks post-inoculation. The plants were grown under a 12-hour photoperiod with a 20°C night temperature and 28°C day temperature and irrigated with a dilute nutrient solution as needed to maintain adequate soil moisture. Because these experiments utilized a non-California isolate of the pathogen, the experiments were quarantined and conducted in compliance with federally-mandated biosafety regulations (https://www.aphis.usda.gov/aphis/home/)

### SNP Genotyping

DNA was isolated from newly emerged leaves harvested from field grown plants using a previously described protocol (Pincot et al., 2020). Leaf samples were placed into 1.5 ml tubes or coin envelopes and freeze-dried in a Benchtop Pro (VirTis SP Scientific, Stone Bridge, NY). Approximately 0.2 g of dried leaf tissue/sample was placed into wells of 2.0 ml 96-well deep-well plates. Tissue samples were ground using stainless steel beads in a Mini 1600 (SPEX Sample Prep, Metuchen, NJ). Genomic DNA (gDNA) was extracted from powdered leaf samples using the E-Z 96® Plant DNA Kit (Omega Bio-Tek, Norcross, GA, USA) according to the manufacturer’s instructions. To enhance the DNA quality and yield and reduce polysaccharide carry-through, the protocol was modified by adding Proteinase K to the lysis buffer to a final concentration of 0.2 mg/ml and extending lysis incubation to 45 min. at 65°C. Once the lysate separated from the cellular debris, RNA was removed by adding RNase A. The mixture was incubated at room temperature for 5 min. before a final spin down. To ensure high DNA yields, the sample was incubated at 65°C for 5 min. following the addition of elution buffer. DNA quantification was performed using Quantiflor dye (Promega, Madison, WI) on a Synergy HTX (Biotek, Winooski, VT).

The individuals phenotyped in these studies were genotyped with either 50K or 850K Axiom® SNP arrays (Hardigan et al., 2020). SNP markers on the 50K Axiom array are a subset of those on the 850K Axiom array. The probe DNA sequences for SNP markers on both arrays were previously physically anchored to the Camarosa and Royal Royce reference genomes (Edger et al., 2019; Hardigan et al., 2020, 2021b). The Camarosa genome assembly has been deposited in the Genome Database for the Rosaceae (https://www.rosaceae.org/species/fragaria_x_ananassa/genome_v1.0.a1) and Phytozome (https://phytozome-next.jgi.doe.gov/info/Fxananassa_v1_0_a1). The Royal Royce genome assembly has been deposited in the Genome Database for the Rosaceae (https://www.rosaceae.org/Analysis/12335030) and Phytozome (https://phytozome-next.jgi.doe.gov/info/FxananassaRoyalRoyce_v1_0). The assemblies for each Royal Royce haplotype have been deposited in a Dryad repository (https://doi.org/10.25338/B8TP7G). The physical addresses for the SNP markers are provided in our online resources (https://doi.org/10.25338/B8TP7G). We utilized both reference genomes as needed to cross-check and compare statistical findings and search genome annotations. The results presented in this paper utilized the haplotype-resolved Royal Royce reference genome FaRR1 (Hardigan et al., 2021a) unless otherwise noted. SNP genotypes were called using the Affymetrix Axiom Suite (v1.1.1.66). Samples with call rates exceeding 89-93% were included in genetic analyses.

### Statistical Analyses of Germplasm Screening Experiments

The R package *lme4* was used for linear mixed model (LMM) analyses of the germplasm screening experiments (Bates et al., 2015). LMMs for square lattice experiment designs were analyzed with entries as fixed effects and incomplete blocks, complete blocks, years, entries × years, and residuals as random effects (Hinkelmann and Kempthorne, 1994). LMMs for randomized complete block experiment designs (ignoring incomplete blocks) were analyzed in parallel to estimate the relative efficiency of the square lattice to the randomized complete block experiment designs (Hinkelmann and Kempthorne, 1994). We did not observe an increase in efficiency by using incomplete blocks; hence, the statistics reported throughout this paper were estimated using LMMs for randomized complete block experiment designs (Hinkelmann and Kempthorne, 1994). Estimated marginal means (EMMs) for entries were estimated using the R package *emmeans* (Lenth, 2017, 2021). Variance components for random effects were estimated using REML (Bates et al., 2015). To estimate broad-sense heritability on a clone-mean basis 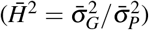, analyses were repeated with entries as random effects, where 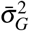 is the among entry variance, 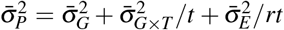 is the phenotypic variance on a clone-mean basis, 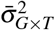 is the entry × year variance, 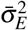 is the residual variance, *t* is the number of years, and *r* is the harmonic mean number of replications. Our experiments were designed with four replications/entry; however, because of the random loss of plants (experimental units), *r* was 3.4 in the 2016-17 field experiment, 4.0 in the 2017-18 field experiment, 4.0 in the 2018-19 growth chamber experiment (for race 1 screening), and 3.7 in the 2018-19 growth chamber experiment (for race 2 screening).

### Genome-Wide Association Study

Genome-wide assocation study (GWAS) analyses were carried out to search for the segregation of loci affecting resistance Fusarium wilt races 1 and 2 among individuals genotyped with either the 50K or 850K Axiom SNP array (Hardigan et al., 2020). GWAS analyses were applied to estimated marginal means (EMMs) for resistance phenotypes using physical positions of SNP markers in the Camarosa and Royal Royce reference genomes (Edger et al., 2019; Hardigan et al., 2021a). SNP marker genotypes were coded 1 for AA homozygotes, 0 for heterozygotes, and -1 for aa homozygotes, where A and a are the two SNP alleles. GWAS analyses were performed using the *GWAS* function in the R package *rrBLUP*. The genomic relationship matrix (GRM, K) was estimated from SNP marker genotypes for each population using the *rrBLUP A*.*mat()* function (VanRaden, 2008; Endelman, 2011). The genetic structure of the GWAS population was investigated using hierarchical clustering and principal components analysis of the GRM as described by Crossa et al. (2014). To correct for population structure and genetic relatedness, a Q + K linear mixed model was used where Q is the population stratification structure matrix and K is the GRM (Yu et al., 2006; Kang et al., 2008). The first three principal components from eigenvalue decomposition of the GRM were incorporated into the Q + K model. Bonferroni-corrected significance thresholds were calculated for testing the hypothesis of the presence or absence of a significant effect. GWAS was repeated in the California population by fitting a SNP marker (AX-184226354) in LD with *FW1* as a fixed effect using the *rrBLUP::GWAS()* function (Endelman, 2011).

### Haplotype Analyses

Thirty-four race 1 resistant (R) and 37 race 1 susceptible (S) individuals from the California population (Hardigan et al. 2020 were selected for haplotype analyses. Using the genotypes for six Axiom array SNP markers in a 414,365-908,422 Mb window on chromosome 2B, the resistant individuals were predicted to be heterozygous or homozygous for the dominant allele (*FW1*), whereas the susceptible individuals were predicted to be homozygous for the recessive allele (*fw1*). Genotype frequencies for the SNP markers used for this analysis are shown in Table S2 (Online Resource 3). These data informed the minimum and maximum haplotype frequencies expected among R and S individuals for SNP markers in LD with the *FW1* locus.

Short-read (150 bp paired-end) DNA sequences were previously produced by whole-genome shotgun sequencing 34 race 1 resistant (R) and 37 race 1 susceptible (S) individuals from the California population (Hardigan et al. 2020; NCBI SRA BioProject ID PRJNA578384 https://www.ncbi.nlm.nih.gov/sra). DNA variants were called by aligning the short-read sequences for these individuals to the haplotype-resolved FaRR1 reference genome (Hardigan et al., 2020). Using only subgenome-specific DNA sequence alignments, we identified a genome-wide set of disomic variant calls that could be used with current phasing and imputation tools. The variant calls were filtered to retain only bi-allelic SNPs in the 0.0-5.0 Mb segment on chromosome 2B (chr_2B:0-5000000). SNPs with minor allele frequencies > 0.05 were retained. Heterozygous variant calls for individual samples with < 3 reads supporting both the reference and alternate alleles were considered ‘missing’ and sites with > 20% missing data after individual sample filtering were dropped. After filtering, 48,491 SNP calls were retained for haplotype phasing in the 0.0-5.0 Mb target segment.

We used pedigree information and the filtered variant calls in VCF format to generate a PLINK (v1.90; http://zzz.bwh.harvard.edu/plink/; Purcell et al. 2007)’.ped’ binary format input file. Variants were phased and imputed from the ‘.ped’ file using SHAPEIT (v2 r900; https://mathgen.stats.ox.ac.uk/genetics_software/shapeit/shapeit.html#home; Delaneau et al. 2013; O’Connell et al. 2014). We ran SHAPEIT in ‘–duohmm’ mode with 15 burn-in, 15 prune, and 30 main steps. The resulting phased haplotypes for the 71 individuals were used to perform a sliding window analysis of local haplotypes in the 0.0-5.0 Mb segment on chromosome 2B. We tested every local window of 15-30 SNPs with a one-SNP offset. Within each 15-30 SNP long window, we evaluated all possible haplotypes against 20 progressively stricter filtering thresholds by increasing the minimum frequency required in R individuals and lowering the maximum allowed frequency in S individuals (Table S3, Online Resource 3). Windows with haplotype frequencies meeting the specified threshold ‘passed’ and were predicted to be more likely to harbor *FW1*. Conversely, windows with haplotype frequencies failing to meet the specified threshold ‘failed and were predicted to be more unlikely to harbor *FW1*. To visualize the distribution of local haplotypes with progressively stricter minimum frequency differences between R and S individuals across the target segment and identify segments predicted to harbor *FW1*, we summed the total number of windows passing a specific filter threshold across the 0.0-5.0 Mb segment, divided the number of passing windows in a 5 kbp range by the total number of passing windows, and repeated this calculation every 5 kbp. This process was repeated for each filter threshold. The value calculated for each 5 kbp region were summed to obtain the ‘cumulative fraction’ of passing windows across all filter thresholds.

Within the susceptible group, we estimated that 100% of the individuals were homozygous for the susceptible allele (*fw1*). Within the resistant group, we estimated that 96% of the individuals were homozygous and 4% were heterozygous for the resistant allele (*FW1*), and deduced that the maximum frequency for the resistant haplotype in the resistance group was > 0.50 (the expectation when every individual is heterozygous). The empirical estimate of the maximum frequency for the resistant haplotype was 0.57, which was consistent with our *a priori* predictions from SNP marker haplotypes (Online Resource 3; Table S2). This value was used as the threshold for the most stringent filter applied in the sliding window analysis.

### Genetic Mapping

SNP markers with ≤ 5% missing data, high quality codominant genotypic clusters, progeny genotypes concordant with parent gentoypes, and non-distorted segregation ratios (*p* < 0.01) were utilized for genetic and quantitative trait locus (QTL) mapping analyses. The linkage phases of the SNP markers were not known *a priori*. The arbitrarily coded SNPs in the original data were in mixed coupling and repulsion linkage phases. The linkage phases of the SNP markers were ascertained using pair-wise recombination frequency estimates, and recoded so that the 100% of the SNP markers were in coupling linkage phase. This was only necessary in the S_1_ populations. The recoded SNP markers were genetically mapped in S_1_ populations using phase-known F_2_ mapping functions. SNP markers were genetically mapped in full-sib populations using phase-known backcross mapping functions from the subset of SNPs that were heterozygous in the resistant parent and homozygous in the susceptible parent. Genetic maps were constructed using the R packages *onemap* and *BatchMap* (Margarido et al., 2007; Schiffthaler et al., 2017) and custom PERL scripts for binning cosegregating SNP markers, calculating pairwise recombination frequencies, and grouping markers using LOD threshold of 10 and maximum recombination frequency threshold of 0.05. The custom PERL scripts are available in the Dryad repository for this paper (https://doi.org/10.25338/B86057). Linkage groups were aligned and assigned to chromosomes using inter-group linkage disequilibrium statistics and percent-identity against the reference genome (Edger et al., 2019; Hardigan et al., 2021a). Marker orders and genetic distances were estimated in parallel using the RECORD algorithm in *Batchmap* with a 25-marker window, window overlap of 15 markers, and ripple window of six markers (Van Os et al., 2005; Schiffthaler et al., 2017). For smaller linkage groups, the window size was reduced incrementally by five to ensure at least two overlapping windows. We used the *checkAlleles, calc*.*errorlod*, and *top*.*errorlod* functions of the R package *qtl* (Lincoln and Lander, 1992; Broman et al., 2003) and custom R scripts to identify and eliminate spurious SNP markers and successively reconstruct linkage groups as described by Phansak et al. (2016). Genetic distances (cM) were estimated from recombination frequencies using the Kosambi mapping function (Kosambi, 1943).

### QTL Mapping

We applied two approaches to scan the genome for the segregation of quantitative trait loci affecting resistance to Fusarium wilt race 1 in S_1_ or full-sib populations genotyped with the 50K Axiom SNP array (Online Resource 4). First, the effects of individual SNP marker loci were estimated using single marker regression as implemented in the R package *qtl* (Broman et al., 2003). The test statistics for each SNP marker locus were plotted against physical positions in the Royal Royce reference genome (Hardigan et al., 2021a). Second, QTL effects were estimated using Haley-Knott interval mapping as implemented in the R package *qtl* (Haley and Knott, 1992; Broman et al., 2003). These analyses used the positions of markers estimated by *de novo* genetic mapping (Online Resources 4-5). Genome-wide significance thresholds (*p* = 0.05) were calculated by permutation testing with 2,000 permutations (Sen and Churchill, 2001). We estimated 95% Bayes confidence intervals for QTL using the *bayes*.*int* function (Broman and Sen, 2009). The percentage of the phenotypic variance (PVE) explained by a SNP marker locus was estimated using the bias-corrected average semivariance method described by Feldmann et al. (2021), where 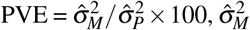 is a bias-corrected REML estimate of the fraction of the genetic variance explained by a SNP marker locus, and 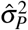 is a REML estimate of the phenotypic variance for resistance to race 1. For statistical analyses of SNP marker loci segregating in S_1_ populations, the additive effect (*â*) was estimated by 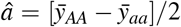, the dominance effect was estimated by 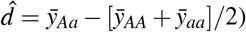, and the degree of dominance was estimated by 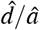, where 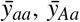, and 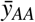 are the respective estimated marginal means (EMMs) for individuals with *aa, Aa*, and *AA* SNP marker genotypes, the *a* was transmitted by the susceptible parent, and the *A* allele was transmitted by the resistant parent. For statistical analyses of SNP marker loci segregating in full-sib populations, effects were estimated the difference between EMMs for heterozygous (*Aa*) and homozygous (*aa*) individuals 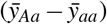.

### KASP Marker Development

Kompetitive allele specific primer (KASP) markers were developed for SNPs predicted to be tightly linked to the Fusarium wilt *R*-genes identified in our studies (Semagn et al. 2014; https://www.biosearchtech.com/support/education/kasp-genotyping-reagents). KASP primers were designed using PolyOligo (Ledda et al. 2020; https://github.com/MirkoLedda/polyoligo) with the Royal Royce reference genome (Hardigan et al., 2021a). We used default PolyOligo design parameters and only tested primers for KASP markers with heuristic quality scores ≥ 7 on a 1 to 10 scale. KASP markers were tested by screening a diverse sample of race 1 resistant and susceptible individuals (*n* = 186) and mapping population progeny. To assess the prediction accuracies of KASP markers, we estimated the concordance between marker genotypes and dominant *R*-gene genotypes inferred from resistance phentoypes (*A*_ for resistant and *aa* for susceptible individuals). KASP-SNP marker genotyping was outsourced to LGC Biosearch Technologies (Hoddesdon, United Kingdom; https://www.biosearchtech.com/support/education/kasp-genotyping-reagents/kasp-overview). The physical locations of SNPs in the Camarosa and Royal Royce reference genomes, oligonucletoide primer sequences, and other supporting data for KASP markers are compiled in Online Resource 5.

## Results

### Resistance to Fusarium Wilt Race 1 is Widespread in the Wild Progenitor Populations and Heirloom and Modern Cultivars of Cultivated Strawberry

Two-thirds of the octoploid strawberry germplasm accessions (ecotypes, cultivars, and other clonally preserved in dividuals) screened for resistance to Fusarium wilt race 1 in the present study (226/344 = 0.66) had disease symptom ratings in the resistant range 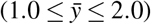, where 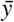 is the estimated marginal mean (EMM) among replicates and years, *y* = 1 plants were symptomless, and *y* = 2 plants were nearly symptomless (Fig. 1; Online Resource 1). The other one-third (118/344 = 0.34) had disease symptom ratings in the moderately to highly susceptible range 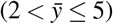. The severity of symptoms (e.g., chlorosis and wilting) increased as 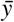 increased on our ordinal scale (plants with scores of five were killed by the pathogen). The race 1 resistance phenotypes observed among resistant and susceptible checks in the present study were consistent with those previously reported (Pincot et al. 2018; Online Resource 1). The repeatability of race 1 resistance phenotypes among clonal replicates of resistant and susceptible checks was 0.81.

**Fig. 1.**
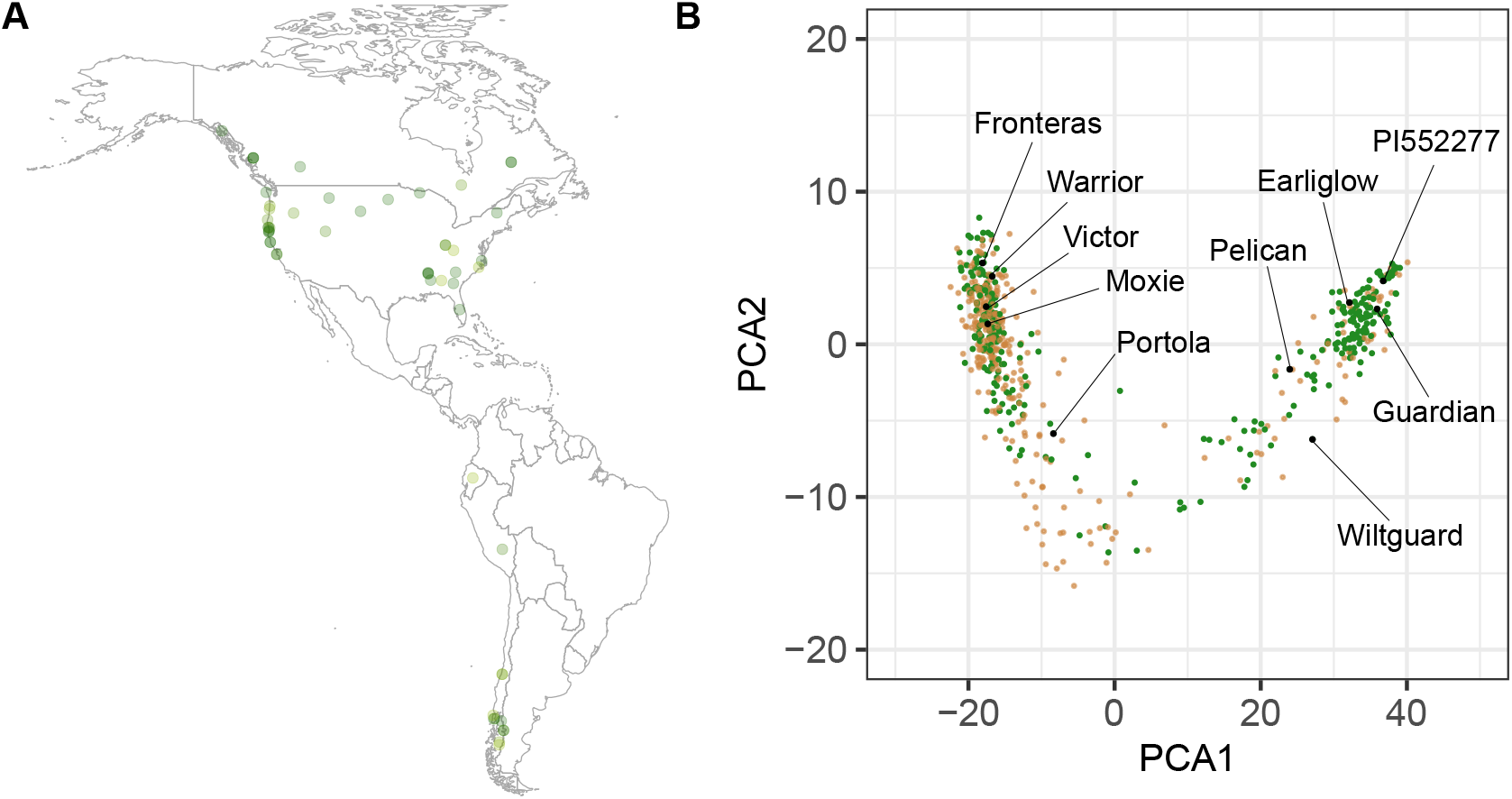
Genetic Diversity of Octoploid Germplasm Accessions Screened for Resistance to Fusarium Wilt Race 1. (A) Geographic distribution (latitude and longitude coordinates) for 27 *F. chiloensis* and 21 *F. virginiana* ecotypes classified as resistant 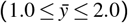 to the AMP132 isolate of *F. oxysporum* f. sp. *fragariae* race 1, where 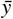 is the estimated marginal mean (EMM) for disease ratings over replications and years (see Online Resource 1 for EMMs and other supporting data). The opaqueness of the points increases as resistance increases (as the EMM decreases). (B) Genetic diversity among 11 *F. chiloensis*, 21 *F. virginiana*, and 608 *F. × ananassa* individuals estimated from the genotypes of 31,212 SNP marker loci assayed with a 50K Axiom SNP array (Hardigan et al., 2020). The first two principal scores from a principal component analysis of the 640 × 640 genomic relationship matrix are displayed with resistant individuals 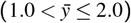 shown in green and susceptible individuals 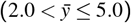 shown in light brown.

One-fourth of the individuals (81/344 = 0.24) screened in the present study were symptomless, classified as highly resistant 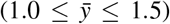, and appeared to be immune to AMP132 infections (Fig. 1; Online Resource 1). This confirmed our suspicion that resistance to race 1 was widespread in natural and domesticated populations of octoploid strawberry. We suspected this because the only ‘non-California’ individuals screened in our previous study (Earliglow and Guardian) were highly resistant to AMP132 infection, had non-*FW1* SNP marker haplotypes, and were presumed to carry novel *R*-genes (Pincot et al., 2018). Moreover, the individuals screened in the present study were more diverse than those previously screened from the California population (Fig. 1; Hardigan et al. 2021b; Pincot et al. 2021).

Slightly more than half of the *F. chiloensis* and *F. virginiana* ecotypes (55/104 = 0.53) screened for resistance to race 1 in the present study were classified as resistant (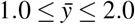; Online Resource 1). We did not observe geographic or phylogenetic trends. Highly resistant ecotypes were found throughout the natural geographic ranges of both species (Fig. 1; Online Resource 1). Eleven out of 62 *F. chiloensis* and 12 out of 40 *F. virginiana* ecotypes were classified as highly resistant 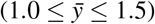. Moreover, highly resistant ecotypes were identified for each of the seven subspecies of *F. chiloensis* and *F. virginiana* we screened other than a single ecotype of *F. virginiana* subsp. *platypetala*, which was nevertheless classified as resistant 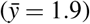. We did not screen ecotypes of *F. chiloensis* subsp. *sandwichensis*, the subspecies found in Hawaii (Staudt, 1989), because none were available when our study was undertaken.

Approximately two-thirds of the *F*. × *ananassa* individuals screened in the present study (160/227 = 0.70) were resistant 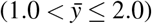 to AMP132 infection (Fig. 2; Online Resource 1). Of these, 20 originated in the California population and were therefore either known or predicted to carry *FW1* (Pincot et al., 2018). The other 140 are heirloom and historically important non-California cultivars or other clonally preserved individuals developed in North America, Europe, and Japan between 1880 and 1987 (Fig. 2; Online Resource 1). Analyses of pedigree records revealed that nearly every one of the resistant non-California cultivars shared resistant common ancestors that dominate the ancestry of modern cultivars (Pincot et al. 2021; Fig. 2); hence, there is a high probability that many of the *R*-alleles found in cultivars worldwide are identical-by-descent.

**Fig. 2.**
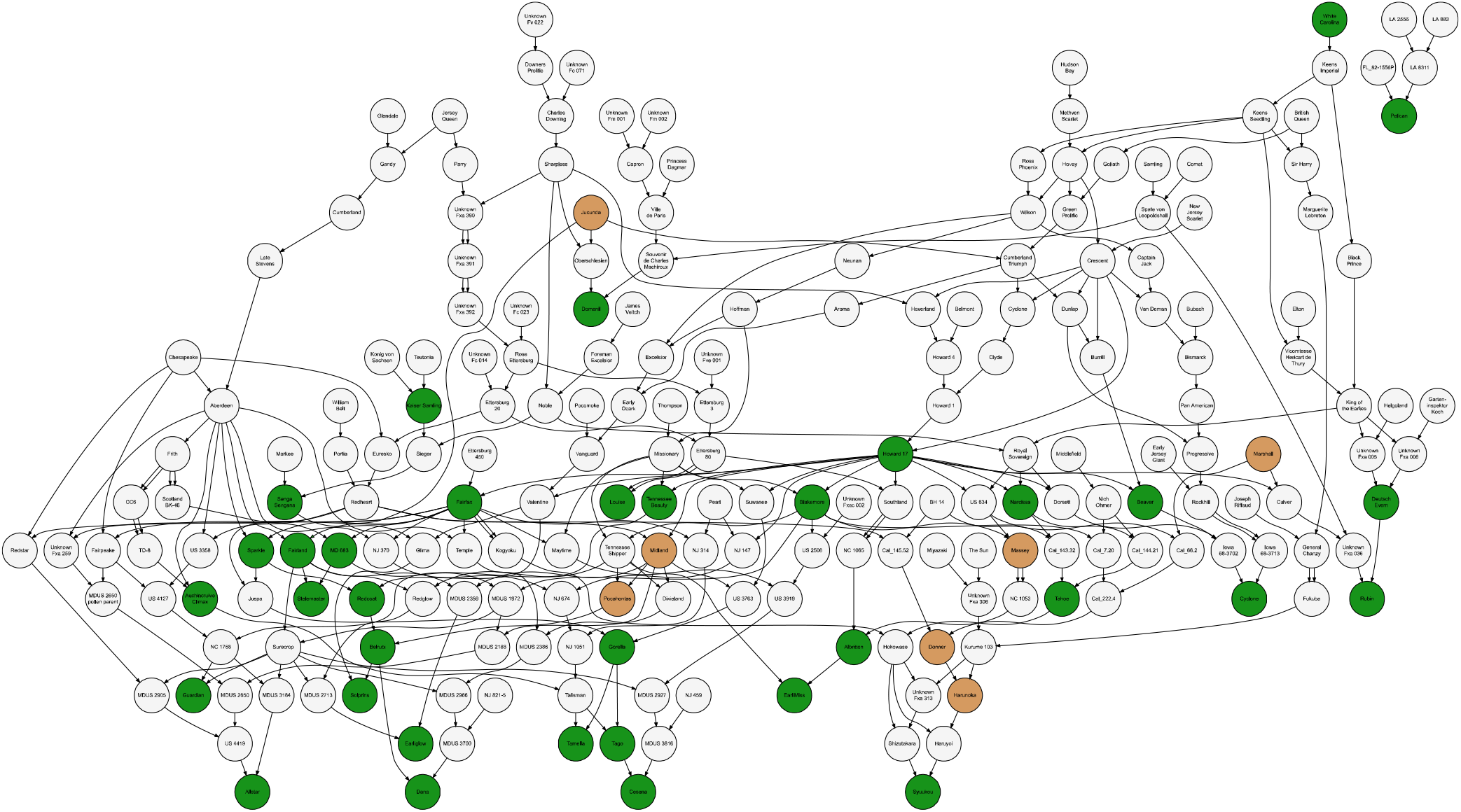
Pedigree Network for Fusarium Wilt Race 1 Resistant Strawberry Germplasm Accessions. Pedigrees are displayed for 117 *F*. × *ananassa* cultivars and other individuals with one or two known parents. The individuals with green or light brown nodes were screened for resistance to the AMP132 isolate of Fusarium wilt race 1. Green nodes identify resistant individuals 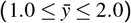 and light brown nodes identify susceptible individuals 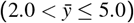, where 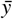 is the estimated marginal mean for resistance to race 1 over replications and years. The race 1 resistance phenotypes of ancestors with light gray nodes are unknown.

### Screening Global Diversity Uncovers Several Sources of Resistance to Fusarium Wilt Race 2

Selection of individuals for constructing the host differential panel (*n* = 144) was informed by insights gained from screening public germplasm collections for resistance to race 1 (*n* = 981 accessions) and from studies of genetic relationships and population structure in strawberry (Hardigan et al., 2020, 2021b; Pincot et al., 2021). The host differential panel was assembled to maximize the probability of differentiating races and identifying sources of resistance to different Fof isolates (Online Resource 1). The phenotypes for resistance to AMP132, MAFF727510, and four other FOF isolates were previously reported for 25 octoploid individuals on the host differential panel: one *F. chiloensis* ecotype, one *F. viriginiana* ecotype, and 23 *F*. × *ananassa* individuals (Pincot et al., 2018; Henry et al., 2021). MAFF727510 is an Fof race 2 isolate found in Japan (Henry et al. 2021).

To broaden insights into the frequency and distribution of race 2 resistance sources, the phenotypes for resistance to MAFF727510 are reported here for an additional 116 individuals: 10 *F. chiloensis* ecotypes (*n* = 10), 16 *F. viriginiana* ecotype, and 93 *F*. × *ananassa* individuals (Online Resource 1). The latter included cultivars and other individuals selected to broadly sample allelic diversity in California and non-California populations worldwide. Similarly, the ecotypes were selected to sample allelic diversity across the natural ranges of *F. chiloensis* and *F. viriginiana*.

The race 1 and 2 resistance phenotypes observed in these studies were highly repeatable: estimates of broad-sense heritability were *Ĥ*^2^ = 0.98 for resistance to AMP132 (race 1) and *Ĥ*^2^ = 0.91 for resistance to MAFF727510 (race 2). Forty-one individuals on the host differential panel (28.5%) were classified as resistant 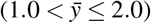 to race 2 (Online Resource 1). Thirty-four of these individuals were symptomless and classified as highly resistant 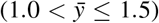, and 34 of the race 2 resistant accessions (87.8%) were resistant to race 1 (Online Resource 1). Of the 78 *F. × ananassa* individuals from the California population, only four (5.1%) were resistant to MAFF727510. Conversely, of the 38 *F*. × *ananassa* individuals from the non-California population, 21 (55%) were resistant to MAFF727510. Slightly more than half of the *F. chiloensis* and *F. viriginiana* ecotypes were resistant to race 2 (16/28 = 0.57), which was comparable to the frequency observed for race 1 resistance (55/104 = 0.53) among 104 ecotypes (Fig. 1; Online Resource 1). Three individuals on the host differential panel (61S016P006, Earlimiss, and Earliglow) were resistant to every Fof isolate tested from California, Japan, Australia, and Spain (Online Resource 1; Henry et al. 2021). According to historical breeding records (Pincot et al., 2021), Royce S. Bringhurt developed 61S016P006 (PI551676), an S_1_ descendant of 43C001P036, by selecting for resistance to Verticillium wilt in Davis, California, nearly a half century before Fusarium wilt was discovered in California (Koike et al., 2009; Koike and Gordon, 2015).

Using the host differential panel as the study population and 50K Axiom SNP array genotypes, we searched the genome for associations between SNP marker loci and race 2 resistance phenotypes. Statistically significant GWAS signals for loci affecting resistance to race 2 were not observed (Online Resources 2 and 3). We repeated this analysis with the host differential panel using race 1 resistance phenotypes and reproduced the strong GWAS signal associated with the segregation of *FW1* in the California population (Online Resources 2 and 3). The absence of a significant GWAS signal for race 2 resistance has several possible explanations. First, resistance to race 2 might not be governed by gene-for-gene resistance. Second, resistance to race 2 could be governed by gene-for-gene resistance but undetectable in a highly diverse population where multiple alleles and loci are segregating and the resistant alleles are uncommon. Those alleles, however, could almost certainly be uncovered and identified by forward genetic analyses of segregating populations developed from crosses between resistant and susceptible parents, as described below for the race 1 resistance genes identified in the present study. Third, the sample size (*n* = 144) may have been insufficient to detect the presence of gene-for-gene resistance to race 2. This seems unlikely because *R*-genes have large effects, and we have consistently observed strong GWAS signals for *FW1* in small samples of California population individuals, including the host differential panel (Online Resources 2 and 3).

### Mendelian Genetic Analyses Uncover the Segregation of Race 1 Resistance Genes in Populations Developed With Ancestrally Diverse Resistant Donors

The sheer numerical abundance of sources of resistance to Fusarium wilt race 1 in strawberry did not shed light on the diversity of *R*-genes that they might carry, if any, or genetic mechanisms underlying resistance (Fig. 1-2). Was resistance to race 1 mediated by dominant *R*-genes? How many unique Fusarium wilt *R*-genes exist in natural and domesticated populations of strawberry? To explore these questions, we developed and undertook genetic analyses of S_1_ populations developed by self-pollinating highly resistant *F*. × *ananassa* heirloom cultivars (Earliglow, Guardian, and Wiltguard) and individuals (61S016P006 and 17C327P010) and full-sib populations developed by crossing highly resistant ecotypes of *F. chiloensis* subsp. *chiloensis* 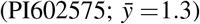, *F. virginiana* subsp. *virginiana* 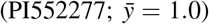, and *F. virginiana* subsp. *grayana* 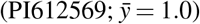 with a highly susceptible *F*. × *ananassa* individual 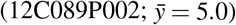 (Table 1).

**Table 1.** Goodness-of-Fit Statistics for Mendelian Genetic Analyses of the Segregation of Fusarium Wilt Race 1 Resistance Genes

| Population | Resistant Parent | Resistant Parent Taxon | Segregation Ratio (R:S)<sup>a</sup> | | $\chi^2$ | $Pr > \chi^2$ |
| --- | --- | --- | --- | --- | --- | --- |
|  |  |  | Observed | Expected |  |  |
| 61S016P006 S<sub>2</sub> | 61S016P006 | *F. × ananassa* | 77 : 0 | - | - | - |
| PI612569 × 12C089P002 | PI612569 | *F. virginiana* subsp. *grayana* | 83 : 0 | - | - | - |
| 12C089P002 × PI602575 | PI602575 | *F. chiloensis* subsp. *chiloensis* | 35 : 41 | 1 : 1 | 0.47 | 0.49 |
| PI552277 × 12C089P002 | PI552277 | *F. virginiana* subsp. *virginiana* | 54 : 57 | 1 : 1 | 0.08 | 0.78 |
| Guardian S<sub>1</sub> | Guardian | *F. × ananassa* | 74 : 25 | 3 : 1 | 0.00 | 0.95 |
| Wiltguard S<sub>1</sub> | Wiltguard | *F. × ananassa* | 72 : 26 | 3 : 1 | 0.12 | 0.73 |
| Earliglow S<sub>1</sub> | Earliglow | *F. × ananassa* | 80 : 5 | 15 : 1 | 0.02 | 0.89 |
| 17C327P010 S<sub>1</sub> | 17C327P010 | *F. × ananassa* | 118 : 8 | 15 : 1 | 0.00 | 0.96 |

<sup>a</sup> The offspring in each segregating population were assigned to resistant (R; $1.0 < y \leq 2.0$ ) and susceptible (S; $2.0 < y \leq 5.0$ ) classes, where $y$ was the visual disease symptom rating on a one to five ordinal scale. The resistant parents were hypothesized to either be homozygous (AA) or heterozygous (Aa) for a partially to completely dominant allele (A), whereas the susceptible parents were hypothesized to be homozygous for a recessive allele (a). Test statistics were estimated using expected ratios for the segregation of either one dominant resistance gene (1:1 for full-sib and 3:1 for S<sub>1</sub> populations) or two unlinked dominant resistance genes with duplicate epistasis (15:1 for S<sub>1</sub> populations).

The phenotypes of offspring in each of the segregating populations we studied spanned the entire range from highly resistant (*y* = 1) to highly susceptible (*y* = 5) with bimodal distributions (Online Resource 3). When individuals within each population were classified as resistant 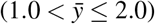 or susceptible 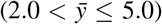 using 2.0 as the cutoff on the disease symptom rating scale, the observed phenotypic ratios perfectly fit the expected phenotypic ratios for the segregation of dominant resistance genes (Table 1). The Guardian and Wiltguard S_1_ and 12C089P002 × PI602575 and PI552277 × 12C089P002 full-sib populations each appeared to segregate for a single dominant resistance gene, whereas the Earliglow and 17C327P010 S_1_ populations appeared to segregate for two dominant genes with duplicate epistasis, where a single dominant allele at either locus was sufficient to confer resistance (Table 1). The statistical inferences were not affected by shifting the cutoff downward to 1.5 or upward to 2.5; hence, we concluded that dominant *R*-genes segregated in these populations (Table 1).

### Fusarium Wilt Resistance Genes are Found on Three Non-Homoeologous Chromosomes

With strong evidence for the segregation of dominant *R*-genes in the Wiltguard and Guardian S_1_ and 12C089P002 × PI602575 and PI552277 × 12C089P002 full-sib populations, we undertook genome-wide searches for associations between SNP marker and causal loci (Table 1; Online Resource 4). The phenotyped individuals from each population were genotyped with the 50K Axiom SNP array (Hardigan et al., 2020), which yielded genome-wide frameworks of SNP markers anchored to physical positions across the 28 chromosomes (Fig. 3; Online Resource 3-4). The average spacing between SNP marker loci ranged from 1.0 to 8.3 cM (Online Resources 3 and 4). The density of SNP marker loci was lower in the parent-specific genetic maps for *F. chiloensis* (PI552277) and *F. virginiana* (PI602575) than for either *F*. × *ananassa* parent.

**Fig. 3.**
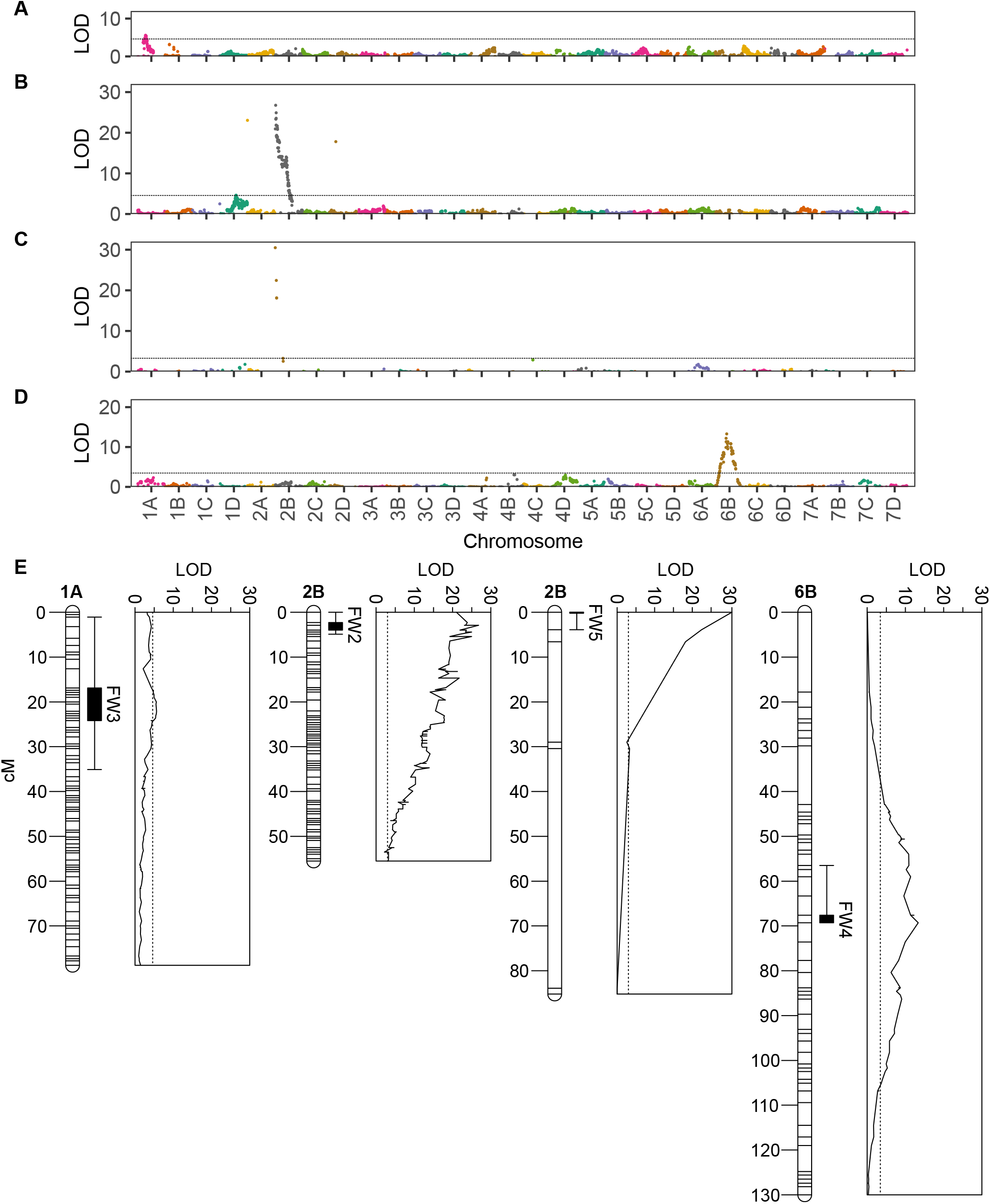
Genome-Wide Search for Associations Between SNPs and Genes Conferring Resistance to Fusarium Wilt Race 1. The upper panels (A-D) display likelihood odds (LODs) for single marker analyses of associations between SNP marker loci and Fusarium wilt race 1 resistance phenotypes in segregating populations genotyped with the 50K Axiom SNP array. LODs are plotted against physical positions of SNP marker loci in the Royal Royce genome. LODs are shown for the Wiltguard S_1_ (A), Guardian S_1_ (B), 12C089P002 × PI602575 full-sib (C), and PI552277 × 12C089P002 full-sib (D) populations. The lower panel (E) displays LODs for analyses of associations between SNP marker loci (plotted along each chromosome) and Fusarium wilt race 1 resistance phenotypes on chromosomes in the Wiltguard S_1_ (*FW3*), Guardian S_1_ (*FW2*), 12C089P002 × PI602575 full-sib (*FW5*), and PI552277 × 12C089P002 full-sib (*FW4*) populations. The dotted lines specify the *p* = 0.05 significance threshold found by permutation testing (*n* = 2, 000). Linkages maps are shown for chromosomes 1A in the Wiltguard S_1_ population, 2B in the Guardian S_1_ and 12C089P002 × PI602575 full-sib populations, and 6B in the PI552277 × 12C089P002 full-sib population. The box and whisker plots display 1-LOD support intervals (solid box) and 95% Bayes confidence intervals (whiskers) for QTL.

We observed a single tightly linked cluster of statistically significant SNP marker loci within each population (Fig. 3A-D) and concluded that they segregated for partially to completely dominant *R*-genes on three nonhomoeologous chromosomes (Fig. 3; Table 2). The putative *R*-genes are hereafter designated *FW2* (inherited from Guardian), *FW3* (inherited from Wiltguard), *FW4* (inherited from PI552277), and *FW5* (inherited from PI602575) (Fig. 3E). *FW2* and *FW5* mapped proximal to *FW1* on chromosome 2B, which suggests that they could be *FW1* alleles or paralogs (Fig. 3E). The genotypic means, effects, and PVE estimates for SNP markers tightly linked with *FW2* and *FW5* were nearly identical to estimates for SNP markers associated with *FW1* in the Fronteras and Portola S_1_ populations (Table 2). The *FW2* allele was nearly completely dominant (*d/a* = 0.85). The additive and dominance effects of the *FW2* locus were 1.5- to 1.9-fold greater than those reported for the *FW1* locus, partly because unfavorable (susceptible) allele homozygotes were more strongly susceptible in the Guardian S_1_ population than in the Fronteras and Portola S_1_ populations. The estimated marginal means (EMMs) for favorable (resistant) allele homozygotes ranged from 1.07 to 1.25 in the three populations (Table 2). The EMM for resistant homozygotes (*FW2FW2* = *AA*) was 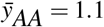, whereas the EMM for susceptible homozygotes (*fw2fw2* = *aa*) was 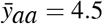. We could not estimate the degree of dominance for *FW5* because the *AA* homozygote was not observed in the full-sib population; however, the EMM for the heterozygote was 1.06, which implies that the *FW5* allele could be completely dominant. We found that a single copy of the *FW2* or *FW5* gene was sufficient to confer resistance to Fusarium wilt race 1.

**Table 2.**
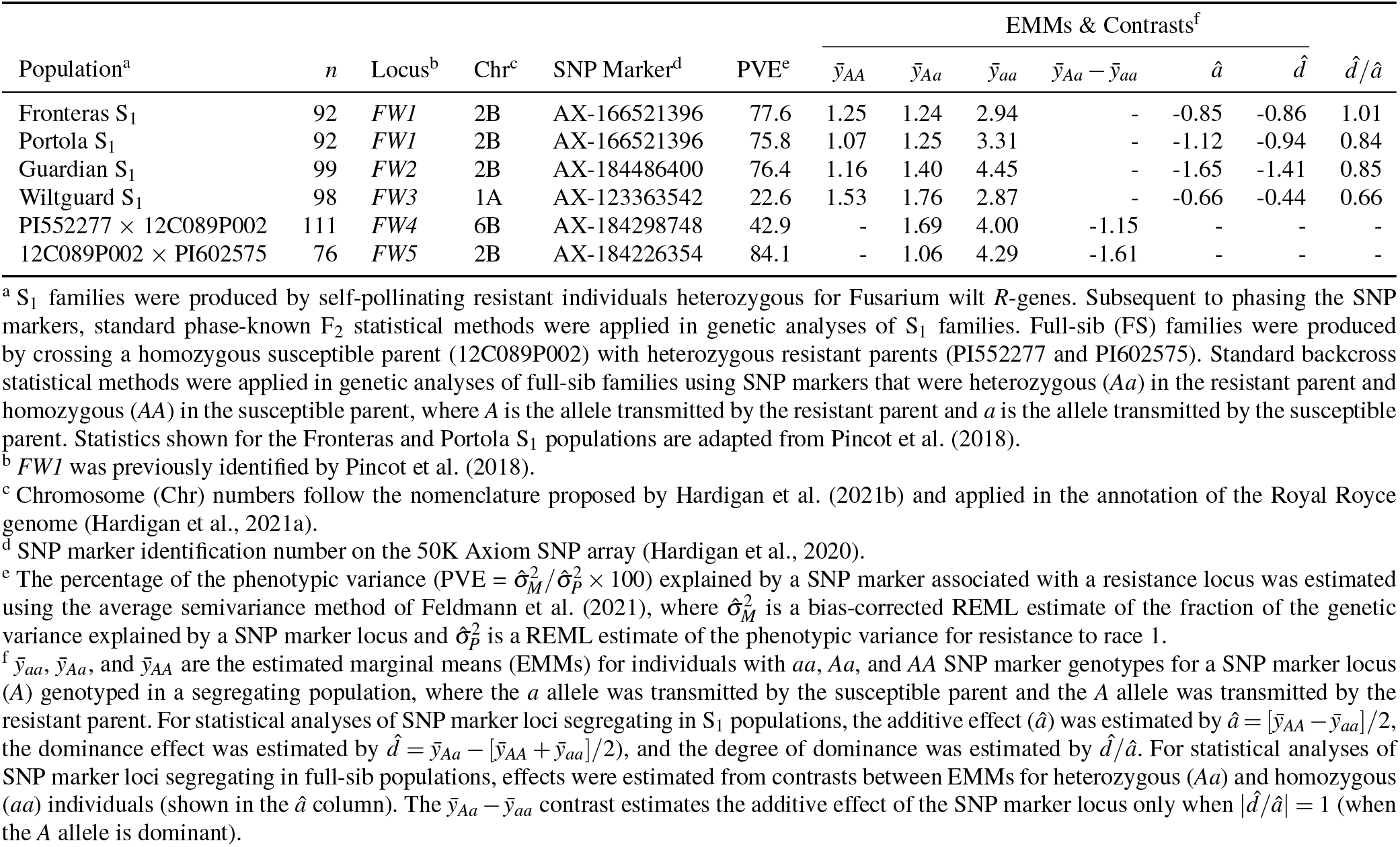
Statistics for SNP Markers Associated With Genes Conferring Resistance to Fusarium Wilt Race 1

Although the statistical evidence for the segregation of a single dominant *R*-gene on chromosome 1A was strong in the Wiltguard S_1_ population, the effects of SNP marker loci associated with *FW3* were weaker than those observed for SNP marker loci associated with *FW1, FW2*, and *FW5* on chromosome 2B (Table 2). The most significant *FW3*-associated SNP marker was AX-123363542 (LOD = 5.6), which only explained 23% of the phenotypic variation for resistance to race 1. Despite this, the EMM for *FW3* homozygotes 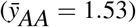 was only slightly greater than the EMMs for *FW1* and *FW2* homozygotes (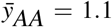 and 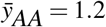, respectively). The *FW4* gene on chromosome 6B appears to be as strong as *FW3* (Table 2). Although the PVE estimate was greater for *FW4* (42.9%) than *FW3* (22.6%), the EMMs for heterozygotes were virtually identical: 1.76 for *FW3* and 1.69 for *FW4*.

### Association Mapping Pinpointed the *FW1* Locus to a Short Haploblock on Chromosome 2B

The *FW1* locus was originally physically mapped using diploid genome-informed GWAS in a closed breeding population of 565 California population individuals genotyped with a SNP array populated with diploid genome-anchored SNPs (Pincot et al., 2018). To revisit our original analyses using octoploid genome-informed GWAS, 356 of these individuals were genotyped with either the 50K or 850K Axiom SNP arrays (Fig. 6). This substantially increased the density and uniformity of SNPs in the *FW1* haploblock and was essential because the probes sequences for the SNP markers on these arrays were anchored *in silico* to octoploid reference genomes developed since the original study was reported (Edger et al., 2019; Hardigan et al., 2020, 2021a,b).

GWAS pinpointed the location of the *FW1* locus to a near-telomeric haploblock on the upper arm of chromosome 2B spanning approximately 3.1 Mb (Fig. 6; Online Resource 2). We confirmed this by developing and genetically mapping KASP markers for SNPs associated with race 1 resistance phenotypes and showing that they were tightly linked with the *FW1* locus and Axiom array SNP marker loci associated with the *FW1* locus on chromosome 2B in the Fronteras and Portola S_1_ populations (Table 2-3; Fig. 6). These KASP-SNP markers explained 76 to 78% of the phenotypic variance and had prediction accuracies ranging from 98.7 to 98.8% in the Fronteras and Portola S_1_ populations and 95.2 to 97.6% in the California population (Tables 2-3; Fig. 4). The 3.1 Mb haploblock was populated with 1,618 SNP markers from the 50K and 850K Axiom SNP arrays, of which 411 were significantly associated with race 1 resistance phenotypes (Fig. 6). SNP markers with the strongest GWAS signals on chromosome 2B were AX-184226354 (0.414 Mb; -log_10_ *p* = 54.6) and AX-184176344 (-log_10_ *p* = 54.6) (Fig. 6; Table 2).

**Fig. 4.**
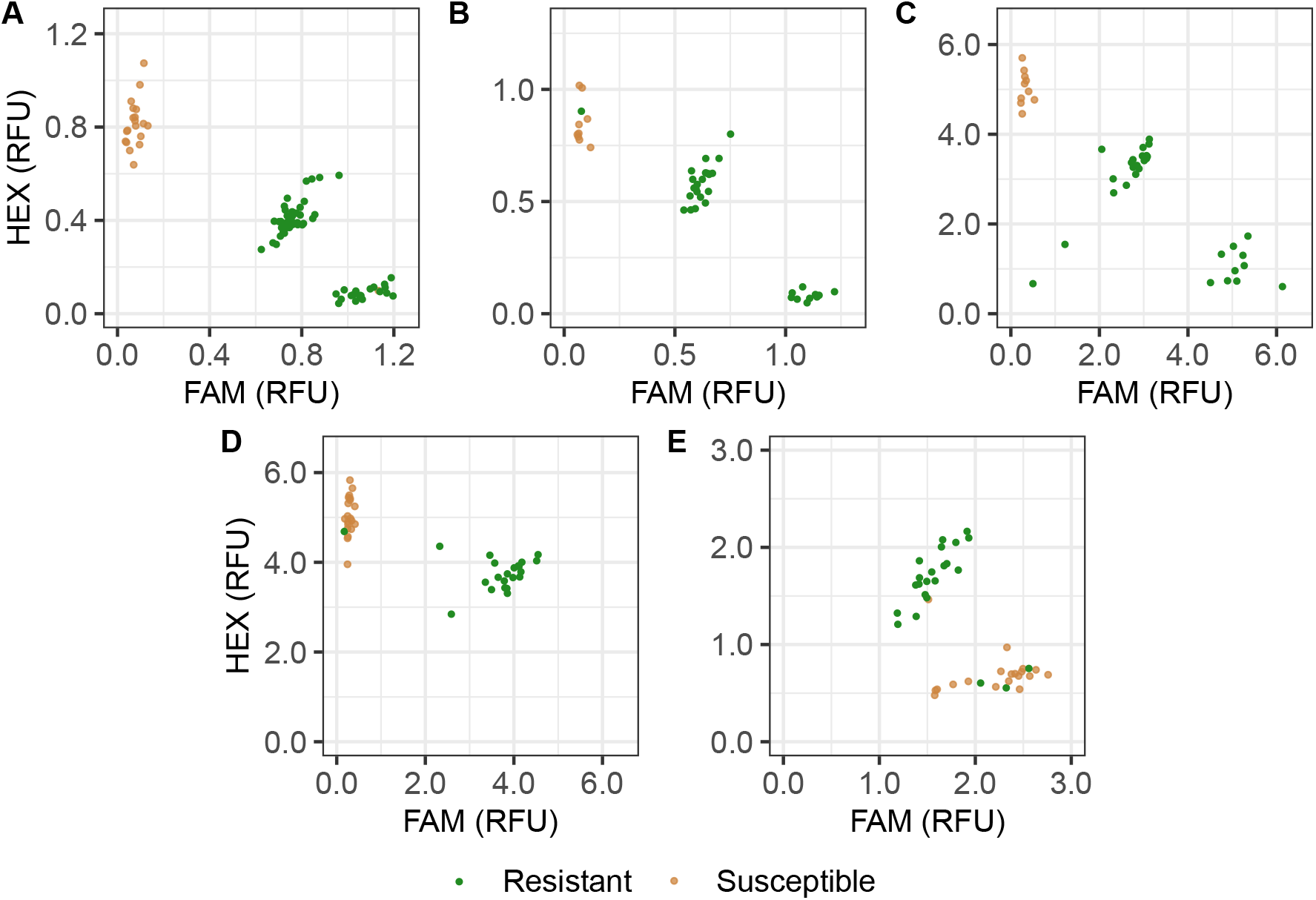
Kompetitive Allele Specific PCR (KASP) Markers for Single Nucleotide Polymorphisms in Linkage Disequilibrium with Fusarium Wilt Resistance (*FW*) Loci in Strawberry. For each KASP marker, individuals labelled *RR* were homozygous for the allele transmitted by the resistant parent, individuals labeled *Rr* were heterozygous, and individuals labeled *rr* were homozygous for the allele transmitted by the susceptible parent. Resistant individuals 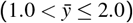 are shown in green, susceptible individuals 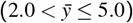 are shown in brown, and 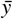 is the estimated marginal mean for disease ratings over replications and years. FAM and HEX signals are reported in relative fluorescence units (RFUs). Fluorescence intensities were normalized using a passive reference dye (ROX). (A) *FW1_K7* KASP marker genotypes observed in the Portola S_1_ (*n* = 40) and the Fronteras S_1_ (*n* = 40) populations for a SNP associated with the *FW1* locus. (B) *FW2_K3* KASP marker genotypes observed in the Guardian S_1_ population (*n* = 40) for a SNP associated with the *FW2* locus. (C) *FW3_K3* KASP marker genotypes observed in the Wiltguard S_1_ population (*n* = 40) for a SNP associated with the *FW3* locus. (D) *FW4_K1* KASP marker genotypes observed targeting *FW4* in the PI552277 × 12C089P002 full-sib population (*n* = 40) for a SNP associated with the *FW4* locus. (E) *FW5_K4* KASP marker genotypes observed in the 12C089P002 × PI602575 full-sib population (*n* = 40) for a SNP associated with the *FW5* locus.

Although GWAS signals were observed between SNP markers and resistance phenotypes on other chromosomes, 84% (68/81) of the 50K and 80% (392/488) of the 850K Axiom array SNP markers with statistically significant GWAS signals were concentrated on chromosome 2B proximal to the previously genetically mapped *FW1* locus (Fig. 5-6; Fig. S3 Online Resource 3; Pincot et al. 2018). The strongest signals, however, were observed for SNP markers AX-89872358 (-log_10_ *p* = 130.9), AX-184098127 (- log_10_ *p* = 122.2), and AX-184513679 (-log_10_ *p* = 117.0), which had previously been assigned *in silico* to chromosome 2D (Hardigan et al. 2020; Online Resource 2). AX-184055143 was the most significant SNP (-log_10_ *p* = 26.6) on chromosome 2D in the 850K analysis (Fig. 6; Fig. S3 Online Resource 3). We show here that the signals observed on other chromosomes in our initial locus-by-locus genome-wide scan were caused by the inaccurate *in silico* assignment of Axiom SNP array probe DNA sequences to physical positions in the octoploid reference genomes (Fig. 5). Three lines of evidence are presented here to substantiate this.

**Fig. 5.**
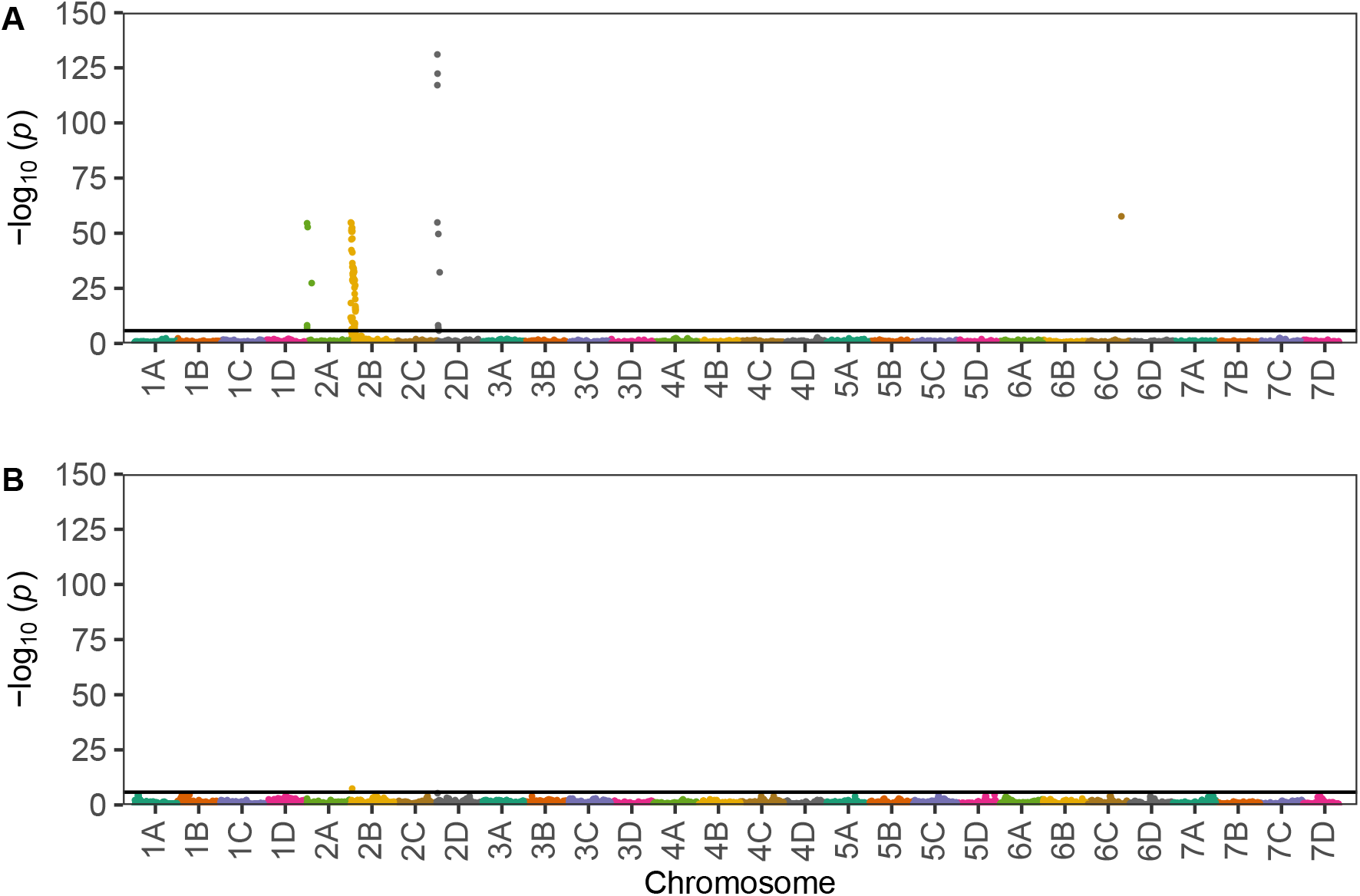
Genome-Wide Associations Between SNP Markers and Fusarium Wilt Resistance Phentoypes. Manhattan plots displaying associations between SNP markers and Fusarium wilt race 1 resistance phenotypes observed among California population individuals phenotyped for resistance to the AMP132 isolate of *F. oxysporum* f. sp. *fragariae*. (A) The upper Manhattan plot displays statistics estimated from the resistance phenotypes of 302 individuals genotyped with the 50K Axiom SNP array (Hardigan et al., 2020). The SNP markers were anchored *in silico* to the Royal Royce genome (Hardigan et al., 2021a). (B) The lower Manhattan plot displays statistics estimated from the same data by fitting the the AX-184226354 SNP marker from chromosome 2B as a fixed effect. The horizontal lines identify the Bonferroni-corrected significance thresholds for hypothesis testing (*p* = 1.6 × 10^−6^).

**Fig. 6.**
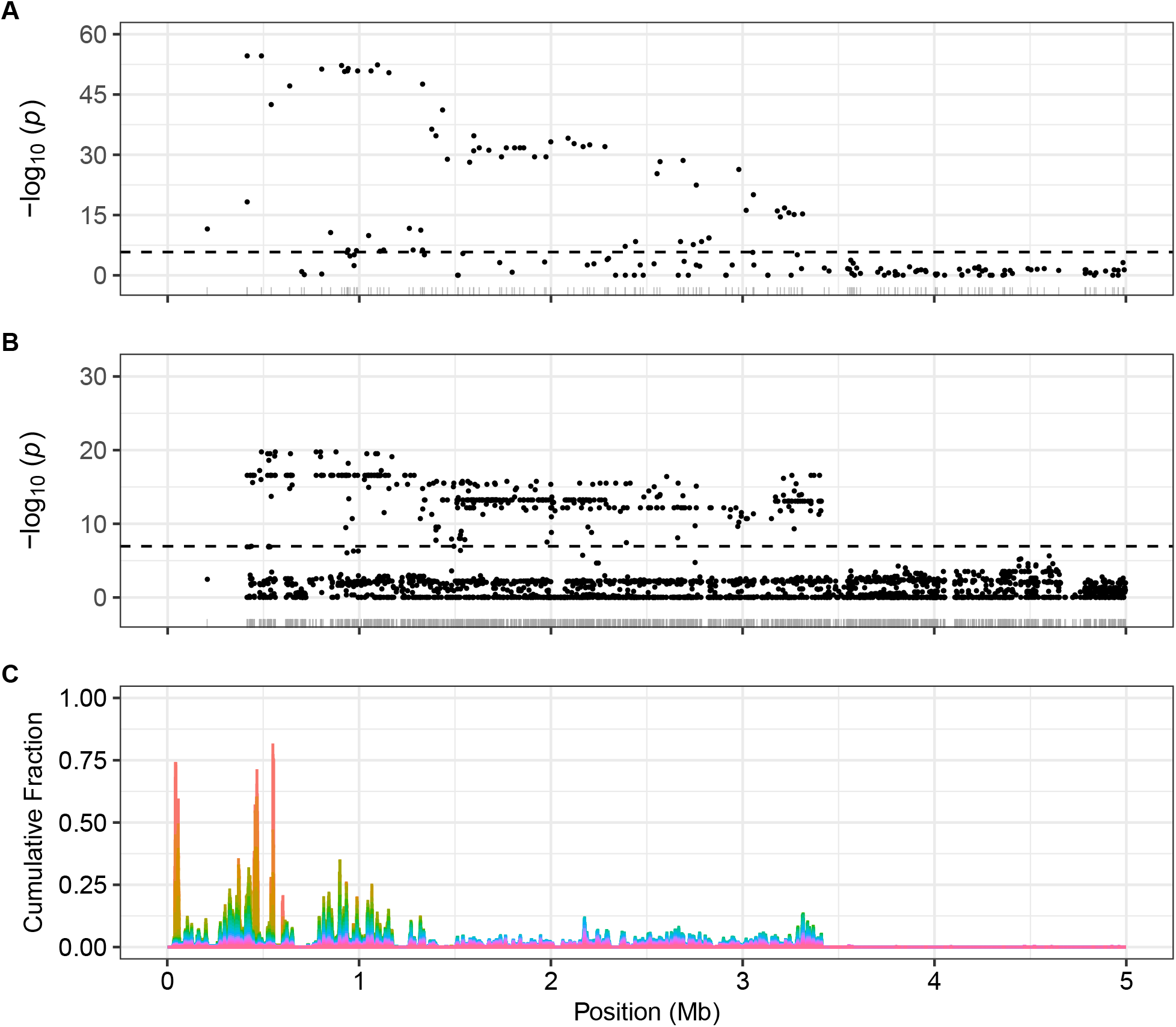
Associations Between SNP Markers and Fusarium Wilt Race 1 Resistance Phentoypes on the Upper Arm of Chromosome 2B. (A) GWAS statistics are shown for the upper 5 Mb haploblock on chromosome 2B from an analysis for race 1 resistance phenotypes among 302 individuals in the California population genotyped with a 50K Axiom SNP array. The individuals in this study were previously phenotyped for resistance using the AMP132 race 1 isolate of *F. oxysporum* f. sp. *fragariae* and predicted to be segregating for *FW*1 (Pincot et al., 2018). The SNP markers were physically mapped to the Royal Royce genome (Hardigan et al., 2021a). Their positions are shown in the rug plot along the x-axis. (B) GWAS statistics are shown from an identical analysis of 54 previously phenotyped individuals in the California population. These individuals were genotyped with an 850K Axiom SNP array. (C) Haplotype analysis of DNA variants between 34 race 1 resistant and 37 race 1 susceptible individuals from the California population. Resistant individuals were predicted to be either homozygous or heterozygous for the dominant (resistant) *FW1* allele, whereas susceptible individuals were predicted to be homozygous for the recessive (susceptible) *FW1* allele. Haplotypes were analyzed in sliding windows of 15-30 SNPs with a 1-SNP offset. The cumulative fractions were estimated by applying 20 progressively stricter filters to identify maximum haplotype frequency differences between resistant and susceptible groups and counting the number of SNP haplotypes in 5 kbp windows. Twenty within group haplotype frequency difference filters were applied to produce the stacked bar plot shown. The colors of each stack correspond to the stringency of the haplotype frequency filters (see Materials & Methods and Table S3 in Online Resource 3).

First, GWAS was repeated by fitting AX-184226354 as a fixed effect, then rescanning the genome for significant GWAS signals. AX-184226354 is a SNP marker on chromosome 2B in strong LD with the *FW1* locus (Table 2-3). The AX-184226354-corrected GWAS completely eliminated GWAS signals on other chromosomes (Fig. 5; Fig. S3 Online Resource 3). To bolster this, we developed KASP markers for the A/G SNP targeted by AX-184055143 (FW1_K1) and T/G SNP targeted by AX-184513679 (FW1_K2); however, neither produced clean codominant genotypic clusters. Nevertheless, we suspect that the probe DNA sequences for these SNP markers reside on chromosome 2B. Second, 16 SNP markers with significant GWAS signals previously assigned *in silico* to chromosomes other than 2B were genetically mapped in present or previous studies (Online Resource 4; Hardigan et al. 2020), and 14 of the 16 were shown to reside on chromosome 2B (83.3%). Third, genome-wide QTL analyses in the Fronteras and Portola S_1_ populations only uncovered statistically significant signals on chromosome 2B (Table 2; Pincot et al. 2018).

A certain percentage of putative off-target signals are expected in GWAS analyses because a certain percentage of the short 71-nt DNA probe sequences for Axiom array SNP markers cannot be unambiguously assigned *in silico* to physical positions in the reference genome, e.g., Hardigan et al. (2020) estimated that approximately 74% of QC-passing 850K Axiom SNP array probes could be assigned to the correct homoeolog in the ‘Camarosa’ genome. That percentage was virtually identical to the percentage of Axiom array SNPs with significant GWAS signals on chromosomes other than 2B in our analyses (Online Resources 2 and 4). This is an important caution for applying GWAS in species with complex repetitive DNA landscapes, especially outbred (heterozygous) species with whole genome duplications where homoeologous DNA variation complicates the physical assignment of short DNA sequences to subgenomes. The assignment of highly accurate long-read DNA sequences, by contrast, is straightforward (Hardigan et al., 2021a). The octoploid genome-informed GWAS analyses described here were initiated in 2017 immediately after we assembled the Camarosa reference genome (Edger et al., 2019), and are the first ever applied in octoploid strawberry, here shown against the highly contiguous haplotype-phased Royal Royce reference genome (Fig. 5). We have since greatly expanded our understanding of the complexity of the octoploid genome, built superior haplotype-phased genome assemblies (Hardigan et al., 2021a), and learned that the challenges associated with differentiating homologous and homoeologous DNA variation are surprisingly minimal in strawberry (Hardigan et al., 2020, 2021b). Our analyses show that the ‘off-target’ DNA marker assignment problem that arises with SNP array-facilitated GWAS can often be solved in polyploid species by fitting DNA markers associated with the target locus as fixed effects, which is effectively equivalent to fitting a multilocus genetic model in a QTL mapping or candidate gene analysis study using mixed linear models (Feldmann et al., 2021).

### Analyses of Phased Haplotypes Saturated the Target Segment With Physically Mapped SNPs and Further Pinpointed the *FW1* Locus

GWAS with the FaRR1 reference genome and 50K and 850K Axiom SNP array genotypes revealed that a short segment (0.0-0.5 Mb) near the upper telomere on chromosome 2B was sparsely populated with SNP markers (Fig. 6). The strongest GWAS signal was close to the border of that segment; hence, we concluded that *FW1* might reside upstream of 0.5 Mb in the SNP sparse segment. To saturate this segment with DNA markers and search for associations between SNPs and *FW1* in the 0.0-5.0 Mb haploblock, short-read DNA sequences for 34 race 1 resistant and 37 race 1 susceptible individuals were aligned to the FaRR1 reference genome (Fig. 6). Using SNP markers tightly linked to the *FW1* locus (chr_2B:414365-908422; FaRR1), we predicted that 100% of the susceptible individuals were homozygous for the recessive allele (*fw1*) and that 96% of the resistant individuals were heterozygous and 4% of the resistant individuals were homozygous for the dominant allele (*FW1*) (Table S2, Online Resource 3). After stringent filtering, 48,491 SNPs were called in the 0.0-5.0 Mb haploblock (1 SNP/103 bp; Fig. 6). SNP haplotypes were phased and imputed using PLINK and SHAPEIT. We searched for SNP haplotypes associated with *FW1* resistance phenotypes by comparing haplotype frequencies between *FW1* resistant and susceptible individuals for 15-30 consecutively phased SNPs in sliding windows with a one SNP offset. The maximum haplotype frequency observed among resistant individuals was 0.5735, whereas the minimum haplotype frequency observed among susceptible individuals was 0.0135 (Table S3; Online Resource 3). These estimates were closely aligned with our *a priori* predictions from SNP marker haplotypes (Table S2; Online Resource 3). Three strong signals were observed between 40,000 and 540,000 bp. The strongest signal was observed between 517,947 to 521,932 bp (Fig. 6; chr_2B:517947-521932) and further narrowed our search for candidate genes for *FW1*.

### Pathogen Defense Genes Associated With Fusarium Wilt Resistance Loci

With the genomic locations of *FW1, FW2*, and *FW5* narrowed to a short haploblock on chromosome 2B (Fig. 6), we searched annotations in the Royal Royce reference genome (FaRR1; Hardigan et al. 2021a) to identify genes encoding proteins known to play an important role in race-specific disease resistance via pathogen recognition and activation of defense responses, e.g., pathogen-associated molecular pattern (PAMP)-triggered immunity or effector triggered immunity (ETI) (Hammond-Kosack and Jones, 1996, 1997; Jones et al., 2016; Zhang and Coaker, 2017; Lolle et al., 2020). Nine of 1,208 annotated genes found in the 0.0-5.0 Mb haploblock on chromosome 2B encode proteins with known *R*-gene domains and functions (Table 4). Four of the nine were found in the 0.0-1.1 Mb haploblock predicted to harbor *FW1*. These included one coiled-coil NLR (CNL) encoding gene (517,947-521,932 bp), one receptor-like kinase (RLK) encoding gene (539,000-554,000 bp), and two tightly linked Toll-interleukin 1 receptor NLR (TNL) encoding genes (1,176,817-1,197,734 bp) (Table 4). Hence, the most promising candidate genes for *FW1* encode CNL and RLK proteins.

The approximate 95% Bayes confidence interval for the genomic location of *FW4* on chromosome 6B (13.8 to 16.3 Mb) was fairly wide and consequently harbored 197 annotated genes in the Royal Royce reference genome (Table 4). Similar to the story for the *R*-gene loci found on chromosome 2B, nine of the 197 annotated genes are predicted to encode *R*-proteins that mediate gene-for-gene resistance in plants (Hammond-Kosack and Jones, 1996; Jones et al., 2016; Zhang and Coaker, 2017; Lolle et al., 2020). These included multiple NBS-LRR *R*-proteins (Table 4). Finally, the approximate 95% Bayes confidence interval for the genomic location of *FW3* on chromosome 1A (4.8 to 8.1 Mb) was slightly wider than that observed for the other mapped loci because the effect of the locus was weaker. There were 535 annotated genes within that interval, of which seven were predicted to encode NBS-LRR or other *R*-proteins (Table 4). This was the locus with the weakest support for the segregation of a race-specific *R*-gene; however, as noted earlier, homozygous resistant (*FW3FW3*) offspring in the Wiltguard S_1_ population were highly resistant (EMM = 1.53). Hence, even if *FW3* does not encode a race-specific *R*-protein, this locus merits further study, in part because the favorable allele (*FW3*) can be deployed and pyramided to strengthen and increase the durability of resistance to Fusarium wilt.

### High-Throughput SNP Genotyping Assays for Marker-Assisted Selection of Fusarium Wilt Resistance Genes

To accelerate the introduction and selection of Fusarium wilt resistance genes in breeding programs, we developed a collection of high-throughput Kompetitive Allele Specific PCR (KASP) markers for SNPs in linkage disequilibrium with *FW1*-*FW5* (Table 3; Fig. 4). Collectively, 25 KASP markers were designed for the five loci using PolyOligo 1.0 (https://github.com/MirkoLedda/polyoligo). The genotypic clusters for 17 of these were codominant (non-overlapping), co-segregated with the predicted resistance loci, and were robust and reliable when tested on diverse germplasm accessions (Fig. 4; Online Resource 5). For each target locus, at least one KASP-SNP marker had a prediction accuracy in the 98-100% range when tested in the original populations where they were discovered (Table 3; Fig. 4). To further gauge their accuracy when applied in diverse germplasm, they were genotyped on 78 California and 66 non-California individuals, mostly cultivars (Online Resources 1 and 5). Because the casual genes and mutations underlying *FW1*-*FW5* are not known, the SNPs we targeted are highly population specific (Table 3). They are strongly predictive when applied in populations where specific genes are known to be segregating and moderately predictive when assayed among random samples of individuals because of recombination (LD decay) between the SNP markers and unknown causal mutations.

**Table 3.** Genomic Locations and Prediction Accuracy Statistics for KASP-SNP Markers Associated With Genes Conferring Resistance to Fusarium Wilt Race 1

| KASP Marker Name<sup>a</sup> | Locus | Chr<sup>b</sup> | FaCA1 Genome Position (bp)<sup>c</sup> | FaRR1 Genome Position (bp)<sup>d</sup> | SNP (R/S)<sup>e</sup> | Axiom Array SNP Marker | Discovery Population Prediction Accuracy (%)<sup>f</sup> | CA Population Prediction Accuracy (%)<sup>g</sup> | Non-CA Population Prediction Accuracy (%)<sup>h</sup> |
| --- | --- | --- | --- | --- | --- | --- | --- | --- | --- |
| FW1_K7 | *FW1* | 2B | 20,343 | 432,840 | T/G | AX-184585165 | 98.8 | 97.5 | 35.9 |
| FW1_K6 | *FW1* | 2B | 31,131 | 443,570 | T/G | AX-184950786 | 98.7 | 97.5 | 24.6 |
| FW1_K3 | *FW1* | 2B | 373,024 | 804,139 | G/T | AX-184624236 | 98.8 | 95.0 | 43.8 |
| FW2_K3 | *FW2* | 2B | 505,395 | 951,437 | C/A | AX-184495646 | 97.5 | 62.8 | 75.8 |
| FW2_K4 | *FW2* | 2B | 1,174,541 | NA | T/C | AX-184456942 | 95.0 | 94.0 | 72.3 |
| FW3_K4 | *FW3* | 1A | NA | 6,781,226 | A/G | AX-166511067 | 97.1 | 41.8 | 67.2 |
| FW3_K1 | *FW3* | 1A | NA | 6,878,550 | A/T | AX-123363542 | 100.0 | 45.1 | 35.6 |
| FW3_K3 | *FW3* | 1A | 6,547,713 | 7,028,803 | C/T | AX-184165918 | 100.0 | 34.6 | 38.1 |
| FW3_K5 | *FW3* | 1A | 6,574,293 | 7,054,363 | A/G | AX-184204436 | 100.0 | 45.0 | 36.5 |
| FW4_K2 | *FW4* | 6B | 22,154,455 | 13,987,603 | A/G | AX-184854002 | 97.4 | 56.4 | 77.4 |
| FW4_K1 | *FW4* | 6B | 21,416,161 | 14,800,900 | T/G | AX-184298748 | 97.5 | 22.9 | 76.9 |
| FW4_K5 | *FW4* | 6B | 22,542,704 | 15,815,205 | C/T | AX-184366576 | 95.0 | 56.8 | 51.7 |
| FW4_K3 | *FW4* | 6B | 19,888,153 | 16,348,827 | T/C | AX-184030353 | 95.0 | 26.8 | 76.9 |
| FW5_K4 | *FW5* | 2B | 1,491,222 | 1,732,659 | G/T | AX-184348754 | 90.0 | 45.7 | 70.3 |

<sup>a</sup> KASP-SNP markers are identified by the Fusarium wilt resistance locus and an alphanumeric suffix starting with a K and ending with an integer.
<sup>b</sup> Chromosome (Chr) numbers follow the nomenclature proposed by Hardigan et al. (2021b).
<sup>c</sup> Physical position of the SNP in the Camarosa reference genome (Edger et al., 2019).
<sup>d</sup> Physical position of the SNP in the Royal Royce reference genome (Hardigan et al., 2021a).
<sup>e</sup> The SNP allele transmitted by the resistant (R) parent is shown to the left, whereas the SNP allele transmitted by the susceptible (S) parent is shown to the right of the slash.
<sup>f</sup> Prediction accuracy statistics for KASP-SNP markers genotyped in a random sample of 40 individuals within each of the original segregating populations developed to discover the Fusarium wilt resistance loci. The prediction accuracy estimates shown here are the frequencies with which SNP marker genotypes correctly identified resistant and susceptible individuals in a particular study population.
<sup>g</sup> Prediction accuracy statistics for KASP-SNP markers genotyped among 86 individuals in the California population.
<sup>h</sup> Prediction accuracy statistics for KASP-SNP markers genotyped among 37 non-California *F. × ananassa* cultivars, 12 *F. chiloensis* ecotypes, and 17 *F. virginiana* ecotypes.

**Table 4.** Pathogen Defense Genes in Linkage Disequilibrium With Fusarium Wilt Resistance Loci

| Locus | Annotation<sup>a</sup> | Chr<sup>b</sup> | Position<sup>c</sup> |  | AT_Gene<sup>d</sup> | Domain<sup>e</sup> | Protein Family<sup>f</sup> |
| --- | --- | --- | --- | --- | --- | --- | --- |
|  |  |  | Start | Stop |  |  |  |
| *FW1* | Fxa2Bg200055 | 2B | 517,947 | 521,932 | AT3G07040.1 | NB-ARC | Disease resistance protein (CC-NBS-LRR) |
| *FW1* | NA | 2B | 539,000 | 554,000 | AT4G23160 | LRR | Receptor-like kinase |
| *FW1* | Fxa2Bg200141 | 2B | 1,176,817 | 1,177,501 | AT5G36930.1 | NB-ARC | Disease resistance protein (TIR-NBS-LRR) |
| *FW1* | Fxa2Bg200143 | 2B | 1,191,345 | 1,197,734 | AT4G12010.1 | NB-ARC | Disease resistance protein (TIR-NBS-LRR) |
| *FW1* | Fxa2Bg200175 | 2B | 1,417,268 | 1,420,357 | AT1G09970.1 | LRR | Receptor-like protein |
| *FW1* | Fxa2Bg200271 | 2B | 2,147,842 | 2,150,028 | AT3G43740.1 | LRR | Receptor-like protein |
| *FW1* | Fxa2Bg200289 | 2B | 2,230,151 | 2,231,286 | AT3G07040.1 | NB-ARC | Disease resistance protein (CC-NBS-LRR) |
| *FW1* | Fxa2Bg200404 | 2B | 3,254,299 | 3,257,893 | AT4G34220.1 | LRR | Receptor-like kinase |
| *FW1* | Fxa2Bg200412 | 2B | 3,333,973 | 3,335,142 | AT2G15320.1 | LRR | LRR family protein |
| *FW3* | Fxa1Ag101064 | 1A | 6,037,746 | 6,048,242 | AT5G17680.2 | NB-ARC | Disease resistance protein (TIR-NBS-LRR) |
| *FW3* | Fxa1Ag101128 | 1A | 6,419,064 | 6,423,914 | AT5G66330.1 | LRR | Receptor-like kinase |
| *FW3* | Fxa1Ag101177 | 1A | 6,682,945 | 6,685,470 | AT3G50950.1 | NB-ARC | Disease resistance protein (CC-NBS-LRR) |
| *FW3* | Fxa1Ag101301 | 1A | 7,378,435 | 7,381,245 | AT3G59410.1 | Kinase | Protein kinase |
| *FW3* | Fxa1Ag101393 | 1A | 7,976,850 | 8,032,406 | AT5G66900.1 | NB-ARC | Disease resistance protein (CC-NBS-LRR) |
| *FW3* | Fxa1Ag101404 | 1A | 8,089,827 | 8,100,427 | AT5G66900.1 | NB-ARC | Disease resistance protein (CC-NBS-LRR) |
| *FW4* | Fxa6Bg102048 | 6B | 15,186,176 | 15,189,317 | AT3G57830.1 | LRR | Receptor-like kinase |
| *FW4* | Fxa6Bg102076 | 6B | 15,542,705 | 15,545,819 | AT5G06940.1 | LRR | Receptor-like kinase |
| *FW4* | Fxa6Bg102106 | 6B | 15,792,021 | 15,794,795 | AT2G41820.1 | LRR | Receptor-like kinase |

<sup>a</sup> Annotated gene name in the Royal Royce genome (Hardigan et al., 2021a).
<sup>b</sup> Chromosome (Chr) numbers follow the nomenclature proposed by Hardigan et al. (2021b).
<sup>c</sup> Physical position of the annotated gene in the Royal Royce genome.
<sup>d</sup> The Arabidopsis Information Resource gene identification number (https://www.arabidopsis.org/index.jsp).
<sup>e</sup> Domain architecture abbreviations are nucleotide binding-ARC (NB-ARC) and leucine rich repeat (LRR).
<sup>f</sup> The protein family abbreviations are coiled-coil nucleotide binding site-LRR (CC-NBS-LRR) and Toll-interleukin 1 receptor NBS-LRR (TIR-NBS-LRR).

## Discussion

The deployment of Fusarium wilt resistant cultivars has become critical in California since the early 2000s when outbreaks of the disease were first reported (Koike et al., 2009; Koike and Gordon, 2015). This disease has rapidly spread and become one of the most common biotic causes of plant death and yield losses in California, the source of 88-91% of the strawberries produced in the US (http://www.agmrc.org/commodities-products/fruits/strawberries; https://www.nass.usda.gov/). The scope of the problem was initially unclear, as were the solutions, because the resistance phenotypes of commercially important cultivars, genetic mechanisms underlying resistance, and distribution and race structure of the pathogen were either unknown or uncertain when the disease unexpectedly surfaced in California (Koike and Gordon, 2015; Pincot et al., 2018). A breeding solution instantly emerged with the discovery of *FW1* (Pincot et al., 2018), and was further strengthened with the discovery of additional homologous and non-homoeologous resistance genes in the present study (Fig. 3; Table 2). Genetic and physical mapping of these race-specific *R*-genes has enabled the rapid development and deployment of Fusarium wilt resistant cultivars through marker-assisted selection. The transfer of *R*-genes from race 1 resistant donors to susceptible recipients via MAS has been incredibly fast because the resistant alleles are dominant, found in heirloom or modern cultivars, and identifiable without phenotyping using SNP markers tightly linked to the causal loci (Online Resources 1 and 5; Table 3; Fig. 6).

Once *FW1* was discovered, we knew that we had a robust solution to the race 1 resistance problem; however, we had virtually no knowledge of the diversity of Fusarium wilt *R*-genes in populations of the wild octoploid progenitors and heirloom cultivars of cultivated strawberry that might be needed to cope with pathogen race evolution (Pincot et al., 2018; Henry et al., 2021). We did not purposefully set out to identify redundant *R*-genes but rather to scour global diversity for ancestrally divergent *R*-genes, both to facilitate *R*-gene ‘stacking’ or ‘pyramiding’ (Poland and Rutkoski, 2016; van Wersch et al., 2020) and inform future searches for sources of resistance to as yet unknown races of the pathogen, in addition to assessing the frequency, diversity, and distribution of *R*-genes in the wild and domesticated reservoirs of genetic diversity (Fig. 1; Online Resource 1). Our results paint a promising picture for the identification of genes for resistance to race 2 and other as yet unknown races of the pathogen. As our phenotypic screening studies showed, the frequency of resistance to race 2 was comparable to that observed for race 1 (Online Resource 1; Henry et al. 2021). Similar to our findings for race 1, the sources we identified for resistance to race 2 were symptomless, which suggests that gene-for-gene resistance might underlie their phenotypes. The genetic basis of resistance to race 2 and other races of the pathogen, however, has not yet been elucidated. There is empirical evidence that resistance to Australian isolates of the pathogen might be quantitative (Mori et al., 2005; Paynter et al., 2014). Henry et al. (2021) showed that the non-chlorotic symptom syndrome caused by Australian Fof isolates (wilt-*fragariae*) differs from the chlorotic symptom syndrome caused by California and Japanese Fof isolates (yellows-*fragariae*). Hence, the genetic basis of resistance to the wilt- and yellows-*fragariae* diseases could be markedly different. We identified several strong sources of resistance to Australian and other non-California isolates of the pathogen that should accelerate the discovery of novel race-specific *R*-genes, elucidation of genetic mechanisms, and development of resistant cultivars (Online Resource 1; Henry et al. 2021).

Growing resistant cultivars is indisputably a highly effective and cost-free method for preventing losses to Fusarium wilt race 1 in strawberry (Table 2; Fig. 3). We estimate that approximately two-thirds of the cultivars grown in California since the earliest outbreaks in 2005 were highly susceptible, whereas the other one-third were highly resistant (Pincot et al. 2018; Online Resource 1). Using the race 1 resistance phenotypes observed in our studies and California Strawberry Commission production statistics (https://www.calstrawberry.com/en-us/market-data/acreage-survey), we discovered that susceptible cultivars have been planted on 49-85% of the acreage in California over the last eleven years (2010-2021). That percentage has hovered between 55 and 59% since 2014. Hence, susceptible cultivars continue to be widely planted in California despite incontrovertible evidence that losses to the disease can be prevented by planting cultivars carrying one of the race-specific *R*-genes we identified (Table 2; Fig. 3; Koike and Gordon 2015; Pincot et al. 2018). Over five years of screening plants that were either naturally infected or artificially inoculated with race 1 isolates of the pathogen, we have not observed visible symptoms on cultivars or other germplasm accessions carrying the dominant *FW1* allele (Online Resource 1; Pincot et al. 2018; Henry et al. 2021). Although private sector cultivars were unavailable for inclusion in our studies, the prevalence of *R*-genes in publicly available germplasm collections and shared ancestry of public and private sector cultivars world-wide suggests that the same *R*-genes are widely found in private sector cultivars (Pincot et al., 2021; Hardigan et al., 2021b). We anticipate that production in California will ultimately shift away from susceptible cultivars, particularly as the incidence of the disease increases and yield losses mount (Koike and Gordon, 2015; Henry et al., 2019).

The cloning and characterization of *R*-genes underlying race-specific resistance is important for developing an understanding of their function and interactions with the pathogen and building the foundation needed to engineer resistance through genome editing or other approaches (Chisholm et al., 2006; Lolle et al., 2020; Chiang and Coaker, 2015; Dong and Ronald, 2019; van Wersch et al., 2020). The *I* genes that confer race-specific resistance to *F. oxysporum* f. sp. *lycopersici* in tomato differ in durability and function and provide a model for future studies in strawberry (Bohn and Tucker, 1939; Sela-Buurlage et al., 2001; Houterman et al., 2009; Catanzariti et al., 2015, 2017). The *I-2* gene had a significantly longer life span than the *I* gene, which was defeated in less than a decade subsequent to deployment (Bohn and Tucker, 1939; Alexander, 1945). The durability differences of the tomato *I* genes have been attributed to differences in the dispensability of avirulence genes or mutations in avirulence genes that defeat known *R*-genes (Catanzariti et al., 2015, 2017). The life spans of the race-specific *R*-genes we identified in strawberry are of course unknown; however, the sheer abundance and diversity of race 1 *R*-genes found in the wild relatives predict that sources of resistance to other races of the pathogen can be rapidly identified and deployed (Fig. 1; Table 2; Online Resource 1). The earliest reports of resistance to California isolates of the pathogen emerged when the disease initially surfaced in field experiments in California where well known cultivars were being grown (Koike et al., 2009; Koike and Gordon, 2015). We have since shown that the resistance phenotypes of those cultivars were mediated by *FW1*; hence, the *FW1* gene has endured for at least 16 years to-date. Our search for diverse *R*-genes was partly motivated by the need to prepare for the havoc created by the inevitable evolution and emergence of novel pathogen races and inadvertent introduction of foreign races of the pathogen through infected plants or soil (Gordon, 2017; Henry et al., 2017, 2021).

Our findings suggest that the resistant germplasm accessions identified in the present study carry one or more dominant Fusarium wilt *R*-genes, and that race 1 *R*-genes are found in a wide range of heirloom and modern cultivars (Table 2-1; Fig. 1-2; Online Resource 1). The latter finding further suggests that race 1 resistance genes are found in domesticated populations worldwide, albeit often at low frequency because they have not been consciously selected, e.g., we previously showed that the frequency of *FW1* was 0.16 in the pre-2015 California population and that *FW1* originated in Shasta and other cultivars released in the 1930s (Pincot et al., 2018), 70 to 80 years before Fusarium wilt was first reported in California (Koike et al., 2009; Koike and Gordon, 2015). The phenotypic and pedigree databases we developed should expedite the identification and incorporation of Fusarium wilt *R*-genes into modern cultivars (Online Resource 1; Fig. 2; Pincot et al. 2021). Our data suggest that a certain percentage of modern cultivars are bound to fortuitously carry Fusarium wilt *R*-genes. That was exactly what we discovered in the California population (Pincot et al., 2018).

The race 1 *R*-genes we identified in cultivated strawberry are predicted to be a small sample of those found in the wild reservoir of genetic diversity (Fig. 1; Table 1-2; Online Resource 1). Our analyses of the pedigree records of heirloom and modern cultivars show that those *R*-genes were fortuitously introduced through early founders and survived breeding bottlenecks predating the late twentieth century emergence of this disease in strawberry (Fig. 2; Winks and Williams 1965; Koike et al. 2009; Pincot et al. 2021). Their chance survival in individuals that dominate the ancestry of domesticated populations worldwide is noteworthy because artificial selection for resistance to Fusarium wilt was not knowingly applied anywhere outside of Australia or Japan until 2015 when breeding for resistance to California specific isolates of the pathogen was initiated (Mori et al., 2005; Paynter et al., 2014, 2016; Pincot et al., 2018). This suggests that genes conferring resistance to Fusarium wilt were fairly common in the founders, which is what our data showed—52% of the wild octoploid individuals screened in the present study were highly resistant to race 1 and predicted to carry dominant race 1 *R*-genes (Online Resource 1). Our analyses of pedigree records further suggest that many of the race 1 *R*-genes found in heirloom and modern cultivars have flowed through common ancestors and thus could be identical-by-descent (Fig. 2; Pincot et al. (2021)).

We cast a wide net in our original phenotypic screening experiments because the genetic basis of resistance was unknown before race-specific *R*-genes were discovered (*FW1*-*FW5*), knowledge was lacking to strategically narrow the search, and the frequency of resistance among accessions preserved in public germplasm collections was unknown. Our phenotypic screens were designed by assuming that we might be searching for a needle in a haystack, primarily because phenotypic screens in tomato and other plant species had shown that genes conferring resistance to Fusarium wilt were uncommon or only found in wild relatives, e.g., the Fusarium wilt *R*-genes in cultivated tomato were transferred from wild relatives (Bohn and Tucker, 1939; Alexander, 1945; Sela-Buurlage et al., 2001; Catanzariti et al., 2015, 2017). Although the domestication and breeding histories of tomato and strawberry are quite different, the frequency of Fusarium wilt resistance in wild relatives are similar. Fifty-two to 57% of the individuals we sampled from wild populations of *F. chiloensis* and *F. virginiana* were resistant to races 1 and 2, which are comparable to the percentages reported for race-specific *R*-genes in the wild relatives of tomato (Bohn and Tucker, 1939; Bournival et al., 1990; Bournival and Vallejos, 1991; Sela-Buurlage et al., 2001).

Wild relatives could certainly become an important source of *R*-genes in strawberry breeding going forward; however, the prevalence of *R*-genes in modern cultivars circumvents the need to introduce alleles from wild relatives or wide crosses, which has often been necessary for the development of Fusarium wilt resistant cultivars in tomato, cotton, and other plants (Sela-Buurlage et al., 2001; Ulloa et al., 2013). The wild relatives of many of the agriculturally important species impacted by this pathogen carry chromosome rearrangements or structural DNA variation that impedes gene flow and the recovery of recombinants, e.g., the tomato *I* genes have been introgressed from wild relatives with interspecific structural variation that suppresses recombination and causes the persistence of unfavorable alleles through linkage drag (Scott and Jones, 1989; Sela-Buurlage et al., 2001; Hemming et al., 2004; Takken and Rep, 2010). Importantly for strawberry, the octoploid progenitors are inter-fertile and have highly syntenic genomes with no known or apparent barriers to gene flow or suppressed recombination in wide crosses (Darrow, 1966; Hardigan et al., 2020). This does not eliminate the linkage drag problem altogether but simplifies the challenge of purging unfavorable alleles introduced by exotic donors in wide crosses (Young and Tanksley, 1989; Fulton et al., 2000). Moreover, cultivated strawberry has emerged from only 250 years of domestication in interspecific hybrid populations between wild relatives and thus has not experienced population bottlenecks on a scale similar to wheat, tomato, and other staples that have undergone 7,000 to 10,000 years of domestication (Darrow, 1966; Hardigan et al., 2020; Pincot et al., 2021).

## Acknowledgements

We are grateful to the California growers and industry leaders that put their confidence in us and took the risk of funding this work before we had produced a single data point. Their steadfast support was critical not only for identifying and deploying breeding solutions to the Fusarium wilt problem that has rapidly emerged in California but for empowering the development of state-of-the-art genomic resources for strawberry that made this research possible. We had the honor of working with Dr. Thomas R. Gordon (1951-2021), a renown plant pathologist, world expert on *Fusarium*, and Distinguished Professor Emeritus at the University of California, Davis. Dr. Gordon sadly passed away on June 27, 2021 before the final draft of this paper was completed.

## Author Contributions

DDAP conducted most of the experimental work, organized and designed experiments, performed statistical analyses, and wrote the first draft of the manuscript. AR and NC contributed experiment work. MJF and MAH contributed statistical analyses. PMH designed, organized, and screened germplasm for resistance to race 2. TRG and PMH provided technical advice and isolates of the pathogen for these studies. MVV and GLC assisted with candidate gene identification. GSC contributed to the design of the studies and managed field experiments and plant propagation. SJK designed the studies, contributed to statistical analyses, and supervised DDAP and the other graduate students and post-doctoral researchers that contributed to these studies. All authors reviewed the manuscript and provided suggestions.

## Data Availability

The data for these studies are publicly available in the online resources and a Dryad repository (https://doi.org/10.25338/B86057). Custom scripts developed for genetic mapping and haplotype analyses were deposited in the Dryad repository. Online Resource 1 is an EXCEL database with Fusarium wilt resistance phenotypes (estimated marginal means) and passport data for every individual screened in the present study. Online Resource 2 is an EXCEL database with GWAS statistics estimated using the physical positions of 50K or 850K Axiom SNP markers in the Camarosa and Royal Royce reference genomes. Online Resource 3 contains histograms for race 1 resistance phenotypes observed in segregating populations, LOD plots for QTL analyses, Manhattan plots for GWAS analyses, genetic mapping summary statistics, and additional haplotype analysis information. Online Resource 4 is a database with QTL mapping statistics and genetic positions (cM) and linkage groups for *de novo* genetically mapped 50K Axiom SNP marker loci in the Guardian S_1_, Wiltguard S_1_, PI552277 × 12C089P002, and 12C089P002 × PI602575 mapping populations. Online Resource 5 is an EXCEL database with physical positions, primer sequences, and other data for KASP-SNP markers. Online Resource 6 is an EXCEL database showing SNP marker genotypes in the haploblock on chromosome 2B predicted to harbor *FW1*.

